# Chemical, molecular, and single cell analysis reveal chondroitin sulfate proteoglycan aberrancy in fibrolamellar carcinoma

**DOI:** 10.1101/2021.12.07.471610

**Authors:** Adam B. Francisco, Jine Li, Alaa R. Farghli, Matt Kanke, Bo Shui, Paul D. Soloway, Zhangjie Wang, Lola M. Reid, Jian Liu, Praveen Sethupathy

## Abstract

Fibrolamellar carcinoma (FLC) is an aggressive liver cancer with no effective therapeutic options. The extracellular environment of FLC tumors is poorly characterized and may contribute to cancer growth and/or metastasis. To bridge this knowledge gap, we assessed pathways relevant to proteoglycans, a major component of the extracellular matrix. We first analyzed gene expression data from FLC and non-malignant liver tissue (n=27) to identify changes in glycosaminoglycan (GAG) biosynthesis pathways and found that genes associated with production of chondroitin sulfate, but not other GAGs, are significantly increased by 8-fold. We then implemented a novel LC-MS/MS based method to quantify the abundance of different types of GAGs in patient tumors (n=16) and found that chondroitin sulfate is significantly more abundant in FLC tumors by 6-fold. Upon further analysis of GAG-associated proteins we found that versican (VCAN) expression is significantly up-regulated at the mRNA and protein levels, the latter of which was validated by immunohistochemistry. Finally, we performed single-cell assay for transposon-accessible chromatin-sequencing on FLC tumors (n=3), which revealed for the first time the different cell types in FLC tumors and also showed that VCAN is likely produced not only from FLC tumor epithelial cells but also activated stellate cells. Our results reveal a pathologic aberrancy in chondroitin (but not heparan) sulfate proteoglycans in FLC and highlight a potential role for activated stellate cells.

**Significance:** This study leverages a multi-disciplinary approach, including state-of-the-art chemical analyses and cutting-edge single-cell genomic technologies, to identify for the first time a marked chondroitin sulfate aberrancy in fibrolamellar carcinoma (FLC) that could open novel therapeutic avenues in the future.

## Introduction

Fibrolamellar carcinoma (FLC) is a rare form of liver cancer that predominantly afflicts adolescents and young adults ^(1, 2)^. Patients with FLC lack standard of care, leaving surgical resection as the primary therapeutic option. Unfortunately, often patients are not eligible for surgery due to metastasis at the time of diagnosis. Paramount to improving patient care is the need to identify molecular pathways that are critical to FLC tumor survival and growth.

FLC is characterized by a ∼400kb heterozygous deletion on chromosome 19, which creates the *DNAJB1-PRKACA* (DP) fusion kinase ^(3)^. DP is found in more than 80% of FLC patients ^(4)^ and genome-scale analyses have identified widespread alterations in chromatin activity ^(5)^ leading to dysregulated genes and non-coding RNAs ^(6, 7)^, including microRNAs ^(8, 9)^. These studies have highlighted increased activation of pro-growth pathways ^(5)^ and increased resistance to cell death ^(10)^.

A characteristic histologic feature of FLC tumors is the presence of thick, fibrous bands ^(11)^. This observation brings into focus the likely importance of the extracellular environment to FLC tumor growth and metastasis, as is the case in many other aggressive cancers as well. Surprisingly though, extracellular matrix composition, including proteoglycans in the pericellular tumor microenvironment, has not been investigated in FLC.

Altered extracellular matrix composition has been identified in numerous cancer types ^(12-14)^. These changes include increased deposition of collagens and fibrillins, matrix remodeling enzymes such as matrix metalloproteinases, and elevated abundance of glycosaminoglycans (GAGs) and associated proteoglycans (PGs). GAGs and PGs are of notable interest as they contribute to both the structure and mechanics of the tumor stroma and enhance extracellular signaling by sequestering and concentrating soluble growth factors. Through these mechanisms, GAGs and PGs play an important role in the processes of angiogenesis, proliferation, and migration ultimately promoting tumor metastasis ^(15-18)^.

GAGs are polysaccharides of varying lengths comprised of repeating disaccharide units ^(19)^. The most common GAGs are hyaluronic acid (HA), heparan sulfate (HS), and chondroitin sulfate (CS). HA chains bind to many components of the extracellular matrix, including collagen. Unlike HA, HS and CS, which differ in disaccharide composition and glycosidic bond linkage patterns, are conjugated to specific core proteins, which are then reclassified as proteoglycans ^(20, 21)^. PGs share many general characteristics, including domains that bind soluble growth factors and stimulate cellular receptors. Additionally, individual PGs can vary in length, and the core proteins can be alternatively spliced, all of which can have affect physiological processes ^(22, 23)^. Finally, polysaccharides can undergo extensive sulfation, the effects of which are essential for these functions ^(24)^.

Examples of PGs that have been well-studied in the cancer context include HS proteoglycan, perlecan (HSPG2), and the CS proteoglycan, versican (VCAN). HSPG2 binds multiple fibroblast growth factors (FGFs) and the vascular and endothelial growth factor (VEGF), and is significantly upregulated in many cancers, including melanoma ^(25, 26)^, breast carcinomas ^(27, 28)^, and glioblastoma ^(29)^. Mouse xenograft models of these cancers in which HSPG2 expression has been ablated show reduced tumor volume ^(16, 28, 30)^. Through similar mechanisms, VCAN is known to promote the metastasis of prostate and breast cancer by promoting platelet-derived growth factor (PDGFA) signaling, interacting with selectins, a family of cellular adhesion molecules, and promoting EGFR signaling ^(31)^. The role of proteoglycans in liver cancer, particularly hepatocellular carcinoma (HCC), has been studied extensively and multiple HS proteoglycans have been identified as important contributors to tumor progression ^(32)^. However, it is unknown whether these mechanisms are shared with FLC.

In this study, we sought to bridge the important knowledge gap on GAGs/PGs in FLC. Specifically, through the combined use of RNA-seq, GAG disaccharide quantification by LC-MS/MS, and single-cell assay for transposase-accessible chromatin (scATAC), we quantified HS and CS abundance and interrogated the activity of PGs at single-cell resolution in FLC and non-malignant liver samples. These studies confirmed that FLC tumor cells preferentially produce chondroitin sulfate and that VCAN is one of the primary proteoglycans in FLC. Moreover, we demonstrated that there is more ATAC signal at the VCAN locus in activated hepatic stellate cells than any other cell type.

## Materials and Methods

### Human Samples

Informed consent was obtained from all individuals. Tumor and adjacent non-malignant liver samples were collected from patients with FLC by surgeons in accordance with the Institutional Review Board protocols 1802007780, 1811008421 (Cornell University) and 33970/1 (Fibrolamellar Cancer Foundation) and provided by the Fibrolamellar Cancer Foundation. Patients included male and female subjects and some samples were collected from the same patient. All samples were de-identified before shipment to Cornell.

### PolyA+ RNA library preparation and sequencing

The 27 RNA-seq datasets analyzed in this study were generated previously ^(5).^ Frozen tumors underwent physical dissociation using a polytron PT1200 E homogenizer (Thomas Scientific, Swedesboro, NJ) and total RNA was isolated using the Total RNA Purification Kit (Norgen Biotek, Thorold, ON, Canada) per manufacturer’s instructions. RNA purity was quantified with the Nanodrop 2000 (Thermo Fisher Scientific, Waltham, MA) or Nanodrop One and RNA integrity was quantified with the Agilent 4200 Tapestation (Agilent Technologies, Santa Clara, CA). Libraries were prepared by the Cornell Transcriptional Regulation and Expression (TREx) Facility using the NEBNext Ultra II Directional RNA kit (New England Biolabs, E7760, Ipswich, MA). Sequencing was performed at the Biotechnology Research Center at Cornell University on the NextSeq500 (Illumina, San Diego, CA).

### Quantitative PCR

Reverse Transcription was performed using the High-Capacity RNA-to-cDNA Kit (Thermo Fisher Scientific). Gene and microRNA expression were quantified with TaqMan Expression assays on a CFX96 Touch Real-Time System thermocycler (Bio-Rad, Hercules, CA). Gene expression assays were normalized to the expression of *RPS9*. Individual gene assay IDs: CSGALNACT1 Hs00218054_m1, VCAN Hs0017642_m1, CHST3 Hs01000045_m1, CHST11 Hs00218229_m1, RPS9 Hs02339424_m1. Expression values reported are averaged across at least three (n=3) biological replicates unless otherwise stated in the main text.

### Immunoblot analysis

FLC and NML tissues were lysed in RIPA buffer containing Halt protease and phosphatase inhibitors (Thermo Fisher Scientific) at 4°C. Lysates were incubated for 30 minutes and centrifuged at 14,000 X g for 10 minutes at 4°C. Total protein in the supernatant was quantified using the BCA Protein Assay Kit (Thermo Fisher Scientific). Samples were denatured in NuPAGE LDS Sample Buffer (Thermo Fisher Scientific) containing 5% β-Mercaptoethanol for 10 minutes at 70°C and loaded to a 12% SDS-polyacrylamide gel. After electrophoresis, samples were transferred to polyvinylidene difluoride membrane and blocked in Tris buffered saline containing 0.5% TWEEN20 (TBST) and 3% bovine serum albumin for 1 hour at room temperature. Membranes were probed for anti-versican GAGβ (1:1000 dilution, rabbit source, Millipore AB1033) or anti-vinculin (1:10000 dilution, mouse source VLN01, Thermo Fischer Scientific MA5-11690) overnight at 4°C and then incubated with goat anti-rabbit HRP linked IgG (1:10000, Cell signaling). Membranes were visualized using a ChemiDoc MP (Bio-Rad, Hercules, CA).

### Immunohistofluorescence analysis

FLC and NML tissues were formalin-fixed at the time of surgery, dehydrated in ethanol, and paraffin embedded for tissue sectioning onto glass slides. Tissue was deparaffinized by two incubations in xylene, followed by one incubation in 1:1 xylene:ethanol (three minutes per incubation). Tissue was rehydrated by incubation in decreasing concentrations of ethanol: twice in 100%, 95%, 75%, and 50% (three minutes per incubation) Finally, tissue was incubated in a flushing water bath for fifteen minutes. Tissue was then incubated in methanol for twenty seconds, followed by equilibration in PBS for at least two minutes. Antigen retrieval was performed by incubating for twenty minutes in preheated 10mM sodium citrate (pH6.0) buffer containing 0.05% tween-20 (weight/volume) submerged in a boiling water bath. Tissue in citrate buffer was then removed for the bath and allowed to cool for thirty minutes. Tissue was then incubated in PBS containing 0.03% tween-20 for two minutes followed by blocking in 10% normal goat serum diluted in PBS containing 0.03% tween-20 for one hour. Tissue was then washed in PBS containing 0.03% tween-20 for two minutes following overnight incubation at 4°C with anti-versican GAGβ (1:100 dilution, rabbit source, Millipore AB1033). Tissue was then washed three times in PBS containing 0.03% tween-20 for fifteen minutes followed by secondary antibody staining for one hour at room temperature (anti-rabbit Alexafluor 488, 1:1000 dilution, goat source, Thermo Fischer Scientific A32731). Tissue was washed three times in PBS containing 0.03% tween-20 for fifteen minutes followed by a five-minute incubation with DAPI to counterstain cell nuclei. Excess DAPI was washed out with two incubations in PBS for ten minutes each. Tissue was then dried briefly, a small volume VectaMount (Vector Labs, Burlingame, CA, H-5000) of was added, and a coverslip was mounted. Images were acquired on an Olympus DP80 microscope with the CellSense Dimension software package. All images received equal brightness balancing with ImageJ software.

### Single cell assay for transposon-accessible chromatin

#### Homogenization

We generated 6 libraries from 70 milligrams of non-malignant liver (NML), primary FLC tumor, and metastatic FLC tumor either with or without collagenase treatment. Tissue was first chopped over dry ice and then dissociated with a Dounce homogenizer using a loose pestle in 2 mL of homogenization buffer (1X HB: 320 mM sucrose, 30 mM calcium chloride, 18 mM magnesium acetate, 60 mM tris(hydroxymethy)aminomethane (TRIS) pH 7.8, 0.1 mM ethylenediaminetetraacetic acid (EDTA), 0.1 % v/v nonyl phenopolyethoxylethanol-40 (NP40), 0.1 mM phenylmethylsulfonyl fluoride (PMSF), 1mM β-mercaptoethanol, 10 mg/mL collagenase IV) for 15-20 strokes. The homogenate was incubated for 3 minutes on ice and then mixed by pipetting 10 times. 6 mL of ATAC-RSB wash buffer (1X ATAC-RSB: 10 mM TRIS pH 7.4, 10 mM sodium chloride, 3 mM magnesium chloride) was added and the homogenate was mixed by pipetting 5 times, incubated on ice for 5 min, and centrifuged at 500 x g for 10 min at 4°C, and the supernatant was removed without disrupting pellet. The nuclei were resuspended in 2 mL of ATAC-RSB wash buffer and then dissociated with a Dounce homogenizer using a loose pestle for 5 strokes, and then a tight pestle for 15-20 strokes, and incubated on ice for 3 min. Then 6 mL of ATAC-RSB wash buffer was added mixed by pipetting-10 times and incubated on ice for 3-5 minutes. The suspension was passed through 70µm Cell Strainer, centrifuged at 500 x g for 10 minutes at 4°C, and the supernatant was removed. The pellet was resuspended in 1 mL OMNI-ATAC buffer (1X OMNI-ATAC: 12.5 mM TRIS pH 7.4, 6.25 mM magnesium chloride, 1.25% v/v dimethylformamide (DMF), 0.125% v/v Tween-20, 0.01% v/v digitonin in 0.4X phosphate-buffered saline (PBS)) and passed through a 30µm Cell Strainer. A 10µL aliquot of nuclei was quantified by double staining with 4’,6-diamidino-2-phenylindole (DAPI) (10 µL of a 100 ug/mL solution incubated 5-10 minutes at room temperature) and trypan blue (10 µL of 0.4% v/v solution incubated 5 minutes at room temperature). The final nuclei count was adjusted to 300,000 nuclei/mL.

#### Tn5 storage and transposome assembly

Tn5 transposase (4 µM) is stored at -80°C and diluted for usage by adding 0.8 volume of 100% glycerol. The transposome is assembled by adding 0.11 volume of Tn5 adaptors (25µM stock solution) to Tn5 stock solution. The mixture was incubated at room temperature for 12-24h. The transposome (∼2µM) can be used directly or stored at -20°C.

#### Tagmentation and Sample Processing

Nuclei suspension (8 µL) was distributed onto 96 well plates and 1 µL of each i5 and i7 transposome (final concentration 400 nM), was added to each well, resulting in a unique barcode combination in each nucleus. The tagmentation reaction plate was incubated (30 minutes at 50°C) and the reaction was terminated by adding 10 µL 20mM EDTA (15 minutes at 37°C). Next, 20µL of Sorting buffer (1X SB: 1X PBS, 2mM EDTA, 20ng/mL bovine serum albumin (BSA)) was added to each well and nuclei were re-pooled into a single sample. Intact nuclei were then stained with DRAQ7 for 15 minutes (ABCam, ab109202, Waltham, MA), passed through a 30 µm filter, and reisolated by fluorescent-assisted cell sorting using a FACSMelody instrument (Becton, Dickinson). A 96 well plate was preloaded with 10 µL of modified sorting buffer (1X SEB: 10 mM TRIS pH 8.0, 12 ng/uL BSA, 0.05% v/v sodium dodecyl sulfate (SDS)), 25 nuclei were distributed into each well, and incubated for 10 minutes at 55°C. Nuclei were further solubilized by adding 2.5 µL of 5% v/v Triton-X100. Libraries were amplified 15 cycles while incorporating standard Nextera (Illumina) barcodes (1µL of 25 µM primer i5:i7 in 25 µL PCR reaction per well).

#### PCR cleanup, Size Selection, and Sequencing

All wells were re-pooled and purified with a MinElute PCR purification kit following the manufacturer’s instructions (Qiagen, 28004) and eluted twice with 20µL of the supplied buffer. The 40 µL elution was further purified and size-selected using magnetic solid phase reversible immobilization (SPRI) beads following the manufacturer’s instructions (Beckman Coulter, A63880) and eluted in 20 µL. Libraries were prepared by the Cornell Transcriptional Regulation and Expression (TREx) Facility using the NEBNext Ultra II Directional RNA kit (New England Biolabs, E7760, Ipswich, MA). Sequencing was performed at the Biotechnology Research Center at Cornell University on the NextSeq500 (Illumina, San Diego, CA).

### ArchR Pipeline for single cell ATAC analysis

#### snATAC-seq preprocessing

Bam files are sorted and indexed using samtools and are converted to fragment files using sinto with – use_chrom ““ and –barcode_regex “(?<=_)(.*)(?=_)” parameters. Fragment files are then sorted and finally used to generate tabix files using tabix.

#### snATAC-seq QC and Dimensionality reduction and clustering analysis

The transcription start site (TSS) enrichment score and fragment number of each nucleus is calculated using ArchR v1.0.1. Nuclei with TSS enrichment score less than 3 and fragment number less than 1,000 are removed. Doublet scores were calculated with default parameters.

We preformed iterative latent semantic indexing by using the ‘addIterativeLSI’ function of ArchR. We then used the default harmony algorithm to correct for batch-effects differences and added clusters using the’addClusters’ function.

#### Identification of marker features

We identified cluster markers using the function ‘getMarkerFeatures’ with default parameters and then applied the ‘addImputeWeights’ function to impute the weights of markers. We visualize marker features using the ‘plotEmbedding’ function. To plot browser tracks we used the ‘plotBrowserTrack’ function and arranged track rows from highest to lowest accessibility using the ‘useGroups’ parameters.

#### Identification of cell-types from snATAC-seq data

We used unbiased approaches to assign cell-type identity to clusters. A pairwise comparison of ArchR defined markers and scRNA-seq liver markers revealed high confidence cell-type assignments for clusters 1-3, and 5-12. Additional GO and KEGG analyses confirmed several cluster assignments.

#### Plotting browser tracks

Accessibility of chromatin surrounding genes of interest is plotted using the default ‘plotBrowserTrack’ function.

### RNA-seq bioinformatic analysis

Paired end RNA sequencing reads were aligned to the human genome (hg38) using STAR (v2.4.2a) and reads aligning to the transcriptome were quantified using Salmon (v0.6). Differential expression was determined with DESeq2.0 (v1.3) using a model that accounts for sequencing facility as a covariate.

### Statistical analysis

Statistical comparisons of quantitative PCR and immunoblot results were made using Student’s t-test. Significant differences in gene expression were determined using DESeq2.0. Graphs were generated in the R software package and error bars represent the standard error.

### Preparation of ^13^C-labeled CS disaccharide calibrants

Four ^13^C-labeled CS disaccharides were prepared from three ^13^C-labeled CS 8-mers, including ^13^C-labeled CS-A 8-mer, ^13^C-labeled CS-C 8-mer and ^13^C-labeled CS-E 8-mer, as described in **Supplemental Figure 1A**. The synthesis of 8-mers was completed using the enzymatic approach (33, 34). The only exception from previously published procedures is that a ^13^C-labeled UDP-GlcA was used to replace unlabeled UDP-GlcA during the synthesis to introduce the ^13^C-labeled GlcA residue to the 8-mer products. The synthesis of UDP-[^13^C]GlcA was completed enzymatically from [^13^C]glucose as described previously ^(35, 36)^. The structures of 8-mer products were confirmed by electrospray ionization mass spectrometry.

The 8-mers were then subjected to the digestion of recombinant chondroitin ABCase (*Flavobacterium heparinum*) to yield the disaccharides. The chondroitin ABCase digestion solution contained 3.55 mL 8-mers (2 mg/mL), 400 μL enzymatic buffer (100 mM sodium acetate/2 mM calcium acetate buffer (pH 6.0) containing 0.1 g/L BSA), and 50 μL of recombinant chondroitin ABCase (3 mg/mL). The reaction mixture was incubated at 37 °C overnight. The extent of reaction completion was monitored by the strong anion exchange chromatography on a Pro Pac PA1 column (9×250 mm, Thermo Fisher Scientific) by measuring the absorbance at 232 nm. The purification of ^13^C-labeled CS disaccharides was performed on the Q-Sepharose fast flow column. Mobile phase A was 20 mM NaOAc, pH 5.0 and mobile phase B was 20 mM NaOAc and 1 M NaCl, pH 5.0. The elution gradient with a flow rate of 1 mL/min was used. The absorbance at 232 nm was scanned and recorded. After purification, the disaccharides were desalted on a Sephadex G-10 column. The quantification of ^13^C-labeled disaccharide calibrants was performed based on the standard curve of commercially available native CS disaccharide standards (Iduron, UK).

### Structure analysis of ^13^C-labeled disaccharide calibrants

A strong anion exchange column Pro Pac PA1 (9×250 mm, Thermo Fisher Scientific) was used to determine the purity of ^13^C-labeled disaccharides after purification. Mobile phase A was 3 mM NaH_2_PO_4_, pH 3.0. Mobile phase B was 3 mM NaH_2_PO_4_ and 2 M NaCl, pH 3.0. The gradient was as follows: 0-20 min 0-20% B, 20-65 min 20-95% B, 65-72 min 95% B and 72-75 min 95-100% B with flow rate of 1 mL/min. The UV absorbance at 232 nm was scanned and recorded. Each disaccharide calibrant was eluted as doublet peaks from ProPac PA1 column. Such profiles are typical for the CS disaccharides due to the chemistry of the anomeric carbon. Both -anomer and -anomer are present in the disaccharides.ESI-MS (Thermo Scientific TSQ Fortis) analysis was used to confirm the molecular weight of each ^13^C-labeled disaccharide. The ESI-MS analysis was performed in the negative ion mode and with the following parameters: Negative ion spray voltage at 3.0 kV, sheath gas at 15 Arb, ion transfer tube temp at 320 °C and vaporizer temp at 100 °C. The mass range was set at 200-800. The measured MWs for the ^13^C-labeled CS disaccharide calibrants are: for Di-0S calibrant was 385.2 (Calc MW = 385.1); for Di-4S calibrant was 465.1 (Calc MW = 465.1); for Di-6S calibrant was 465.1 (Calc MW = 465.1); and for Di-4S6S calibrant was 545.2 (Calc MW = 545.0). From our ESI-MS analysis, we did not observe unlabeled CS disaccharides in the calibrants, suggesting that the calibrants had very high isotopic purity that was suited for the quantitative analysis.

### Quantification of the ^13^C-labeled disaccharide calibrants

The CS native disaccharides (Iduron, UK) were dissolved in water and diluted to the concentration of 5, 10, 20, 40 and 80 μg/mL. 50 μL of the diluted CS was injected into HPLC to make the standard curve to quantify the ^13^C-labeled disaccharide calibrants. The stock solutions of four ^13^C-labeled disaccharide calibrants were diluted to 10 or 20 times and then 50 μL was injected to the HPLC analysis. The concentration of the ^13^C-labeled disaccharide calibrants stocks were determined by comparing the peak areas at 232 nm with unlabeled disaccharides.

### Linear dynamic range determination

Individual stock solutions of four CS unlabeled disaccharides (Iduron, UK) were prepared in water at 1 mg/mL. A stock solution of the mixture of the four unlabeled disaccharides, with the final concentration of 0.25 mg/mL for each disaccharide, was obtained by mixing an equal volume of four individual stock solutions. The linear dynamic range of the working solutions was determined by a serial dilution of the mixture stock solution in water to obtain a range of final concentrations (**Supplemental Figure 1B-E**). The ^13^C-labeled CS disaccharide were added to the linear dynamic range working solutions as an internal standard mixture stock solution to the final concentration of 40 g/mL of ^13^C-labeled di-4S, di-6S and di-4S6S respectively and 8 g/mL of di-0S. The linear dynamic range working solutions containing ^13^C-labeled internal standard were freeze-dried and reconstituted in the 20-μL mouse plasma. The reconstituted solutions were filtered by passing through a YM-3KDa spin column (Millipore) and washed twice with deionized water to recover the disaccharides in the eluent. The AMAC (2-aminoacridone)-derivatization of lyophilized disaccharides was carried out by adding 6 μL of 0.1 M AMAC solution in DMSO/glacial acetic acid (17:3, v/v) and incubating at room temperature for 15 min. Then 6 μL of 1 M aqueous sodium cyanoborohydride (freshly prepared) was added to this solution, where AMAC represents 2-aminoacridone purchased from Sigma Aldrich. The reaction mixture was incubated at 45 °C for 2 h. After incubation, the reaction solution was centrifuged to obtain the supernatant that was subjected to the LC-MS/MS analysis.

### LC-MS/MS analysis

The analysis of AMAC-labeled disaccharides was performed on a Vanquish Flex UHPLC System (Thermo Fisher Scientific) coupled with TSQ Fortis triple-quadrupole mass spectrometry as the detector. The C18 column (Agilent InfinityLab Poroshell 120 EC-C18 2.7 μm, 4.6 × 50 mm) was used to separate the AMAC-labeled disaccharides. Mobile phase A was 50 mM ammonium acetate in water. Mobile phase B is methanol. The elution gradient of from 5-45% mobile phase B in 10 min, followed by isocratic 100% mobile phase B in 4 min and then isocratic 5% mobile phase B in 6 min was performed at a flow rate of 0.3 ml/min. On-line triple-quadrupole mass spectrometry operating in the multiple reaction-monitoring (MRM) mode was used as the detector. The ESI-MS analysis was operated in the negative-ion mode using the following parameters: Neg ion spray voltage at 4.0 kV, sheath gas at 45 Arb, aux gas 15 arb, ion transfer tube temp at 320 °C and vaporizer temp at 350 °C. TraceFinder software was applied for data processing. The normalized peak areas of the ^13^C-labeled calibrants were plotted against the concentrations of linear dynamic working solutions.

### Analysis of CS and HS from tissues

CS was extracted from 5 NML and 11 FLC tissues. All tissues were excised, homogenized, and defatted by suspension and vortex in chloroform/methanol mixtures (2:1, 1:1, 1:2 (v/v)). The defatted tissues were dried and weighed to obtain the dry weight. The dried and defatted tissues were digested with Pronase E (10 mg:1 g (w/w), tissue/Pronase E) at 55 °C for 24 h to degrade the proteins. CS was recovered from the digested solution using a DEAE-column. DEAE column mobile phase A was 20 mM Tris, pH 7.5 and 50 mM NaCl, and mobile phase B was 20 mM Tris, pH 7.5 and 1 M NaCl. After loading the digested solution, the column was washed with 10-column volumes of buffer A to discard the contaminants, following by 10 column-volumes of buffer B to elute the CS fraction. The CS eluting from the DEAE column was desalted using an YM-3KDa spin column and washed three times with deionized water to remove salt. A known amount ^13^C-labeled calibrants were added to the digestion solution. The 200 μL of enzymatic buffer (100 mM sodium acetate/2 mM calcium acetate buffer (pH 7.0) containing 0.1 g/L BSA), and the 60 μL of chondroitin ABCase (3 mg/ml) was added to digest the retentate on the filter unit of the YM-3KDa column. The column was incubated at 37 °C overnight. The CS disaccharides and calibrants were recovered by centrifugation, and the filter unit was washed twice with 200 μL of deionized water. The collected filtrates were freeze-dried before the AMAC derivatization. The AMAC label and LC-MS/MS analysis of the collected disaccharides of tissues was performed as described above. The amount of tissue CS was determined by comparing the peak area of native disaccharide to each calibrant.

HS was extract from 2 NML tissues and 4 FLC tissues respectively. The method for the analysis of HS followed the procedures described in a previous publication ^(37)^.

Three CS disaccharides, including Ddi-2S, Ddi-2S6S and Ddi-2S4S, were only performed relative quantitation as the ^13^C-labeled disaccharides were unavailable. Standard curves of these three disaccharides were generated using unlabeled disaccharide standards that were purchased from Iduron.

## Data availability

Previously published RNA-seq data can be downloaded from Gene Expression Omnibus (GEO) using the following GEO accession number: GSE181922. Previously published RNA-seq and ChRO-seq can be downloaded from the European Genome-Phenome Archive (EGA) using the following EGA accession number: EGAS00001004169.

## Results

### Chondroitin but not heparan sulfate biosynthesis genes are increased in fibrolamellar carcinoma

Tissue samples from patients with FLC were acquired at the time of surgical procedures through a collaboration with the Fibrolamellar Cancer Foundation (FCF) and subjected to RNA extraction and messenger RNA sequencing (n=23 tumor samples and n=4 adjacent non-malignant liver (NML) samples), as previously reported ^(5)^. We then performed an analysis of differential gene expression for enzymes related to glycosaminoglycan (GAG) biosynthesis, focusing on hyaluronic acid (HA), heparan sulfate (HS), and chondroitin sulfate (CS).

The expression of hyaluronic acid synthase 1-3 in FLC did not meet our standard threshold of robust expression (>500 normalized counts), and therefore the HA pathway was not considered for further analysis. We then assessed the expression of genes which catalyze the formation of the common tetrasaccharide linker required for HS and CS proteoglycan production (**Figure 1A**). The initial addition of xylose to serine residues in polypeptide chains is catalyzed by xylosyltransferases (*XYLT1* or *XYLT2*) ^(38)^. This reaction is followed first by the addition of two galactose molecules (catalyzed by β1,4-galactosyltransferase-I (*B4GALT7*) and β1,3-galactosyltransferase-I (*B3GALT6*)), and subsequently by the addition of glucuronic acid (catalyzed by β1,3-glucuronyltransferase-I (*B3GALT3*)) ^(39, 40)^. We found that the expression in FLC of the genes that code for these enzymes meets the threshold criteria but is not significantly different relative to NML.

**Figure 1.**
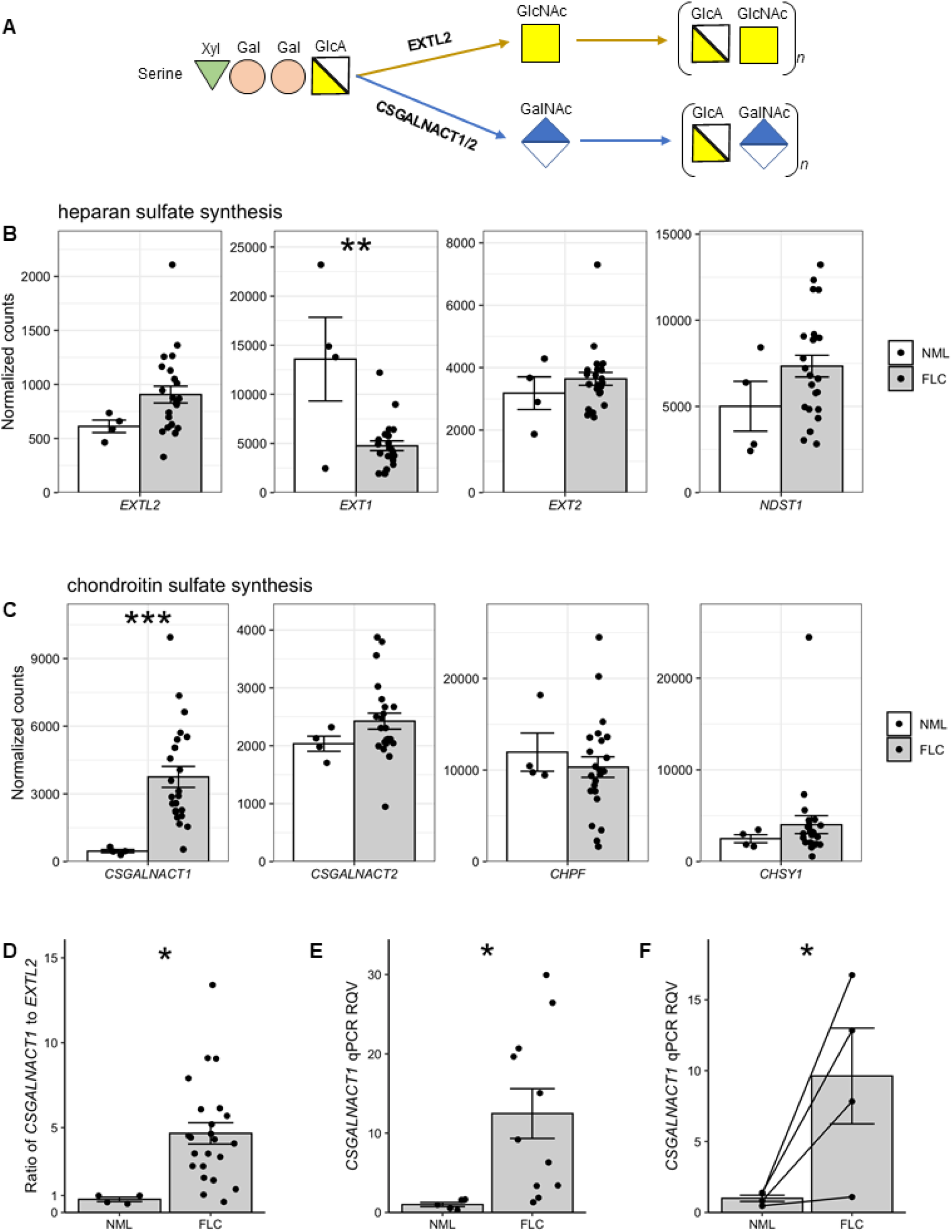
CSGALNACT1 is dramatically up-regulated in FLC. **(A)** A schematic showing the sequence of events leading to heparan and chondroitin synthesis. **(B)** Differential expression of *EXTL2, EXT1, EXT2*, and *NDST1* is shown as normalized counts in FLC (n=23) and NML (n=4). **(C)** Differential expression of *CSGALNACT1, CSGALNACT2, CHPF*, and *CHSY1* is shown as normalized counts in FLC (n=23) and NML (n=4). **(D)** *CSGALNACT1* expression relative to *EXTL2* expression is shown as a ratio of normalized counts in FLC (n=23) and NML (n=4). **(E)** Quantitative PCR showing the relative quantitative value (RQV) of *CSGALNACT1* in a separate cohort of FLC samples (n=11) compared to NML samples (n=5). **(F)** Quantitative PCR showing the relative quantitative value (RQV) of *CSGALNACT1* in a subset of FLC samples (n=4) that have matched NML tissue. The matched NML/FLC samples are indicated with a line linking the two data points. \**P*<0.05, \*\**P*<0.01, \*\*\**P*<0.001, two-tailed Student’s t-test.

We next interrogated those genes encoding enzymes responsible for HS and CS polymerization. HS chain formation begins with the addition of *N*-acetylglucosamine (GlcNAc) to the common linker by *N*-acetylglucosaminyltransferase (*EXTL2*), followed by the addition of glucuronic acid (GlcA) by exostosin glycosyltransferases (*EXT1* and *EXT2*) to create the HS disaccharide ^(20)^ (**Figure 1A**). We found that the expression of *EXTL2* and *EXT2* is unchanged in FLC, and that *EXT1* is significantly decreased (**Figure 1B**). Additionally, HS chains undergo deacetylation and sulfation, catalyzed by the enzyme N-deacetylase and N-sulfotransferase 1 (*NDST1*) ^(41)^, as a critical maturation step and there is no significant change in the expression of this gene in FLC (**Figure 1B**).

CS chain elongation begins with the addition of *N*-acetylgalactosamine (GalNAc) to the common linker by CS GalNAc transferase 1 or 2 (*CSGALNACT1* and *CSGALNACT2*), followed by the addition of GlcA by an enzyme complex containing chondroitin synthase 1 (*CHSY1*) and chondroitin polymerizing factor (CHPF) to create the CS disaccharide ^(42)^ (**Figure 1A**). We found that the expression of *CSGALNACT1* is significantly increased (8-fold, adjusted P=7.6×10^−9^) in FLC compared to NML (**Figure 1C**). The expression levels of *CSGALNACT2, CHPF*, and *CHSY1* are unchanged in FLC; however, *CHPF* is more abundant in FLC than any of the HS polymerizing factors (**Figure 1C**).

Given that HS and CS chains share a common linker, the stoichiometric ratio of *EXTL2* and *CSGALNACT* enzymes is the primary factor determining whether HS or CS chains will be generated ^(43)^. We compared the expression of *CSGALNACT1* to *EXTL2* within each of the FLC and NML samples and found that *CSGALNACT1* is on average ∼4.5 times (P=0.017) more abundant than *EXTL2* (**Figure 1D**) in FLC samples, while being roughly equal in NML samples (0.77 fold). In an independent cohort of patients (FLC n=11 and NML n=4) we confirmed by real-time quantitative PCR (RT-qPCR) that *CSGALNACT1* expression is increased >10-fold (**Figure 1E**). A similar result was obtained when the analysis was restricted only to matched patient samples (**Figure 1F**).

### Chondroitin sulfate chains are aberrantly elevated in fibrolamellar carcinoma

The gene expression analysis is strongly suggestive of increased CS, but not HS, abundance in FLC. To test this hypothesis, we quantified HS and CS abundance in FLC using a novel chemical analytical method. Due to the relatively low abundance of CS from biological tissues, a new quantitative CS analytical method with high sensitivity was developed for this study. Disaccharide analysis is a commonly used approach to analyze the structure of CS polysaccharides. The method involves the degradation of CS polysaccharides into disaccharides using chondroitin ABCase, and the resultant disaccharides were subjected to liquid chromatography coupled with tandem mass spectrometry (LC-MS/MS) analysis (**Figure 2A)**. Furthermore, summing up the amounts of individual disaccharides from the digested CS provides the total amount of CS. To increase the quantitation capability, we employed four ^13^C-labeled CS disaccharide calibrants as internal standards, including di-0S, di-4S, di-6S and di-4S6S (**Supplemental Figure S1**). The ^13^C-labeled CS disaccharide calibrants were obtained from three uniquely designed ^13^C-labeled CS octasaccharides (8-mers) that were synthesized by an enzymatic approach. The di-4S disaccharide is found in CS-A polysaccharide, whereas di-6S and di-4S6S are found in CS-C and CS-E polysaccharides, respectively. The di-0S disaccharide is found in all subtype CS polysaccharides from biological sources. The inclusion of ^13^C-labeled calibrants eliminated batch-to-batch variations, increasing the data consistency (**Supplemental Figure 1B-E**).

**Figure 2.**
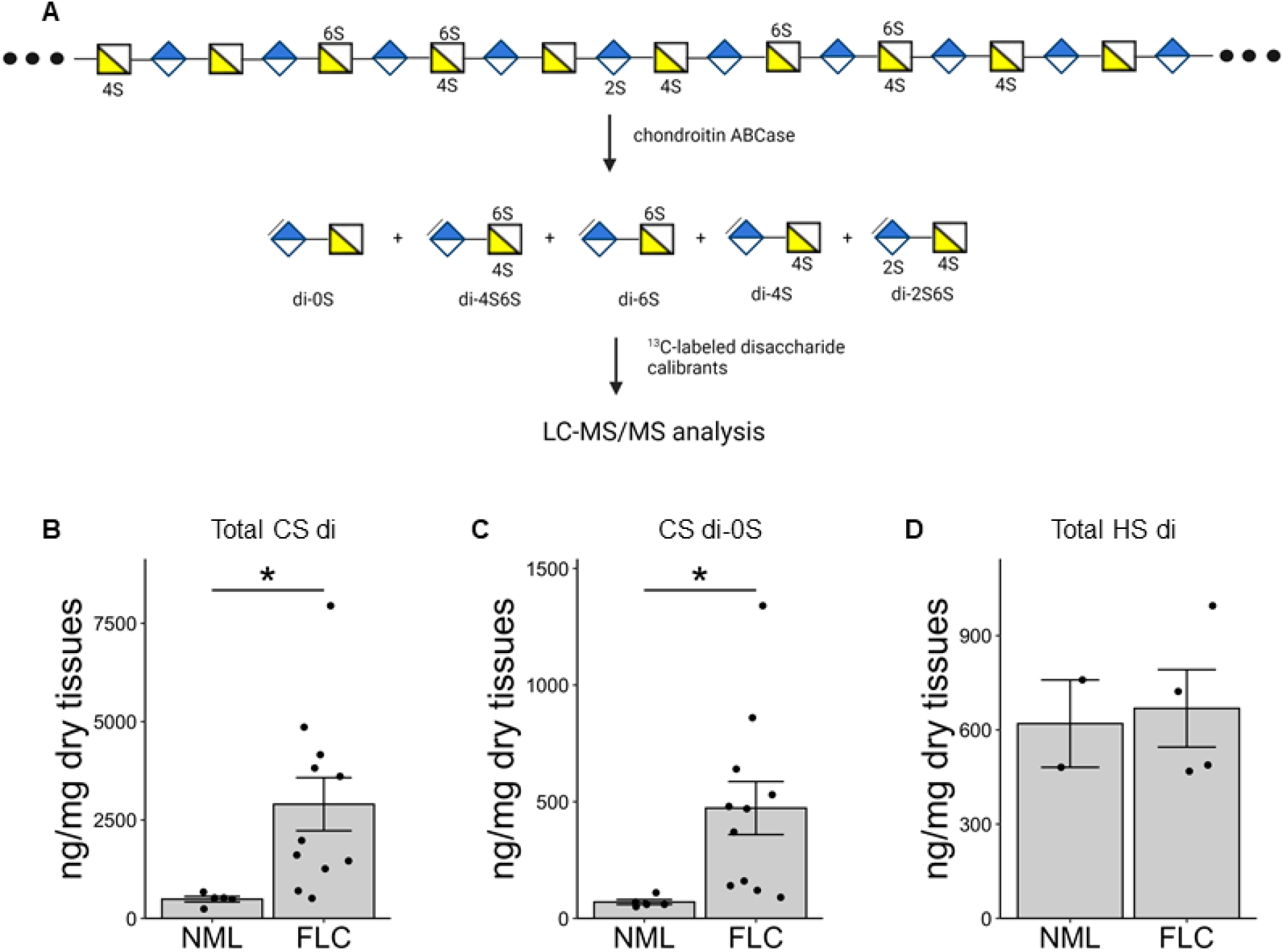
Chondroitin sulfate disaccharide abundance is significantly increased in FLC. **(A)** A schematic diagram detailing disaccharide extraction and identification. **(B)** Nanograms of chondroitin sulfate (CS) per milligram of dry tissue in FLC (n=11) and NML (n=5) tissue. **(C)** Nanograms of non-sulfated chondroitin sulfate (CS di-0S) per milligram of dry tissue in FLC (n=11) and NML (n=5) tissue. **(E)** Nanograms of heparan sulfate (HS) per milligram of dry tissue in FLC (n=4) and NML (n=2) tissue. \**P*<0.05, two-tailed Student’s t-test.

Using this highly sensitive method, we discovered that total CS in FLC tumor tissue compared with non-malignant liver (FLC=11, NML=5) was significantly increased (5.9-fold, P=0.033) (**Figure 2B**) and the amount of non-sulfated CS (di-0S) is also similarly increased (6.7-fold, P=0.035) in FLC (**Figure 2C**). However, there was no difference in total HS in a subset of FLC tumors (FLC=4, NML=2) (**Figure 2D**).

The sulfation of CS chains is mediated by a family of chondroitin sulfotransferase (*CHST*) enzymes that have specificity for particular positions of oxygen on the CS disaccharide. We interrogated the expression of key *CHST* genes in FLC and measured the abundance of sulfated CS. Monosulfation of the 4-OH position of the GalNAc residue (CS di-4S) (**Figure 3A**) is catalyzed by chondroitin 4-*O* sulfotransferase (*CHST11*), and we found that the expression of this gene is significantly increased in FLC (2.6-fold, adjusted P=0.03) (**Figure 3B**). Correspondingly, we also found that CS di-4S is highly elevated in FLC (5.7-fold, P=0.043) (**Figure 3C**). A second common site for monosulfation is at the 6-OH position of the GalNAc residue (CS di-6S) (**Figure 3D**), which is catalyzed by chondroitin 6-*O* sulfotransferase (*CHST3*). We found that although the expression of *CHST3* in FLC is lower than that of *CHST11*, its levels are also significantly increased compared to NML (2.3-fold, adjusted P=0.02 (**Figure 3E**). Likewise, although CS di-6S is not as abundant as CS di-4S, it is similarly increased in FLC (9-fold, P= 0.023) (**Figure 3F**).

**Figure 3.**
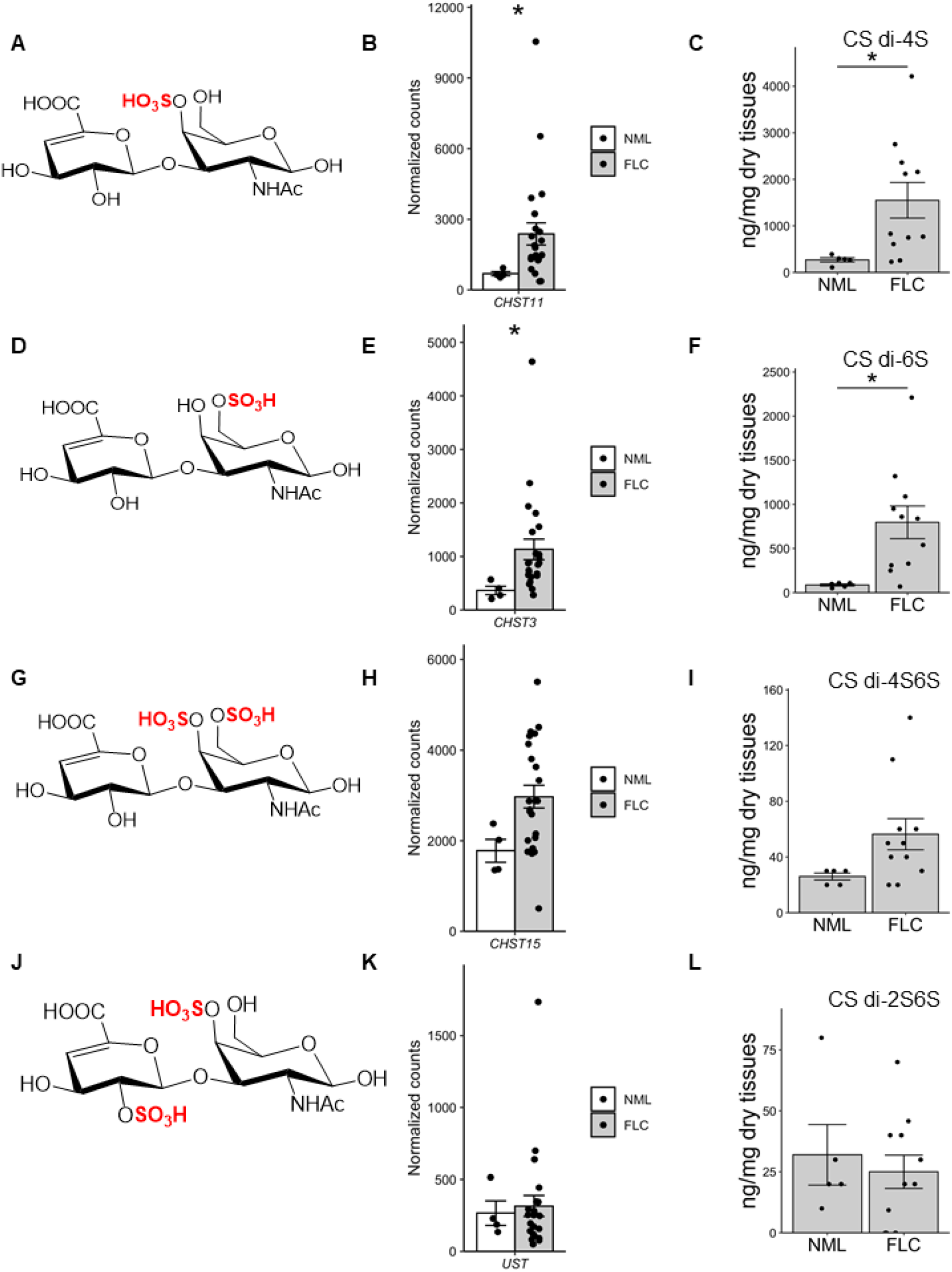
Monosulfated chondroitin sulfate disaccharide abundance is significantly increased in FLC. **(A)** A schematic diagram detailing CS di-4S. **(B)** Differential expression of *CHST11* is shown as normalized counts in FLC (n=23) and NML (n=4). **(C)** Nanograms of CS di-4S per milligram of dry tissue in FLC (n=11) and NML (n=5) tissue. **(D)** A schematic diagram detailing CS di-6S. **(E)** Differential expression of *CHST3* is shown as normalized counts in FLC (n=23) and NML (n=4). **(F)** Nanograms of CS di-6S per milligram of dry tissue in FLC (n=11) and NML (n=5) tissue. **(G)** A schematic diagram detailing CS di-4S6S. **(H)** Differential expression of *CHST15* is shown as normalized counts in FLC (n=23) and NML (n=4). **(I)** Nanograms of CS di-4S6S per milligram of dry tissue in FLC (n=11) and NML (n=5) tissue. **(J)** A schematic diagram detailing CS di-2S4S. **(K)** Differential expression of *UST* is shown as normalized counts in FLC (n=23) and NML (n=4). **(L)** Nanograms of CS di-2S4S per milligram of dry tissue in FLC (n=11) and NML (n=5) tissue. \**P*<0.05, two-tailed Student’s t-test.

CS chains with the disaccharide repeat of di-4S can serve as a substrate for GalNAc 4-sulfate 6-*O*-sulfotransferase (*CHST15*) or uronyl 2-*O*-sulfotransferase (*UST*), respectively, resulting in CS di-4S6S and CS di-2S4S (**Figure 3G, 3J**). We found that the expression of *CHST15* is modestly, but not significantly, increased (**Figure 3H**) and that the expression of *UST* is unchanged (**Figure 3K**) in FLC. Consistent with these findings, we found that the abundance of CS di-4S6S is modestly, but not significantly, increased (**Figure 3I**), and the abundance of CS di-2S4S is unchanged (**Figure 3L**). We performed a similar quantification for HS and found that none of the sulfated subtypes are significantly altered in FLC (**Supplemental Figure 2A-G**), consistent with the finding that total HS abundance is unchanged in FLC (**Figure 1D**). In an independent patient cohort we found by RT-qPCR analysis that the expression of *CHST11* is significantly increased in FLC in all samples (**Supplemental Figure 3A**) as well as in matched samples only (**Supplemental Figure 3B**) and that *CHST3* is trending upward in FLC (**Supplemental Figure 3C and D**). Taken together, these molecular and chemical findings strongly indicate that FLC tumors are marked by aberrant levels of total CS as well as specific sulfated subtypes.

### Versican is the primary chondroitin sulfate-associated protein in fibrolamellar carcinoma

We next assessed changes in the expression of CS associated proteins (CSAPs) in FLC. We identified three CSAPs significantly up-regulated in FLC compared to NML: chondroitin sulfate proteoglycan 8 (*CSPG8*, also known as *CD44*), chondroitin sulfate proteoglycan 4 (*CSPG4*), and versican (*VCAN*). Notably, the fold-change and abundance of *VCAN* in FLC is substantially greater than the other two (*VCAN* ∼10-fold, P=8.5×10^−6^; *CD44* ∼2-fold, P=0.11; *CSPG4* ∼ 4.5-fold, P=0.0016) (**Figure 4A**). We measured *VCAN* by RT-qPCR in an independent FLC patient cohort and observed a significant increase in FLC in all samples (18.5-fold, P=0.0095) (**Figure 4B**) as well as in matched samples only (15.5-fold, P=0.04) (**Figure 4C**). Western blot analysis of three matched FLC/NML pairs confirmed dramatic elevation of VCAN protein in FLC (average ∼200-fold) (**Figure 4D, E**). Specifically, VCAN protein is variable but abundant in the tumor tissue from all three FLC patients, while virtually absent in the adjacent non-malignant samples (**Figure 4D, lower panel**). Finally, we performed immuno-histofluorescent (IHF) staining on two matched FLC/NML pairs of samples and confirmed that VCAN protein is robustly, though non-uniformly, detected only in tumor tissue (**Figure 4F**).

**Figure 4.**
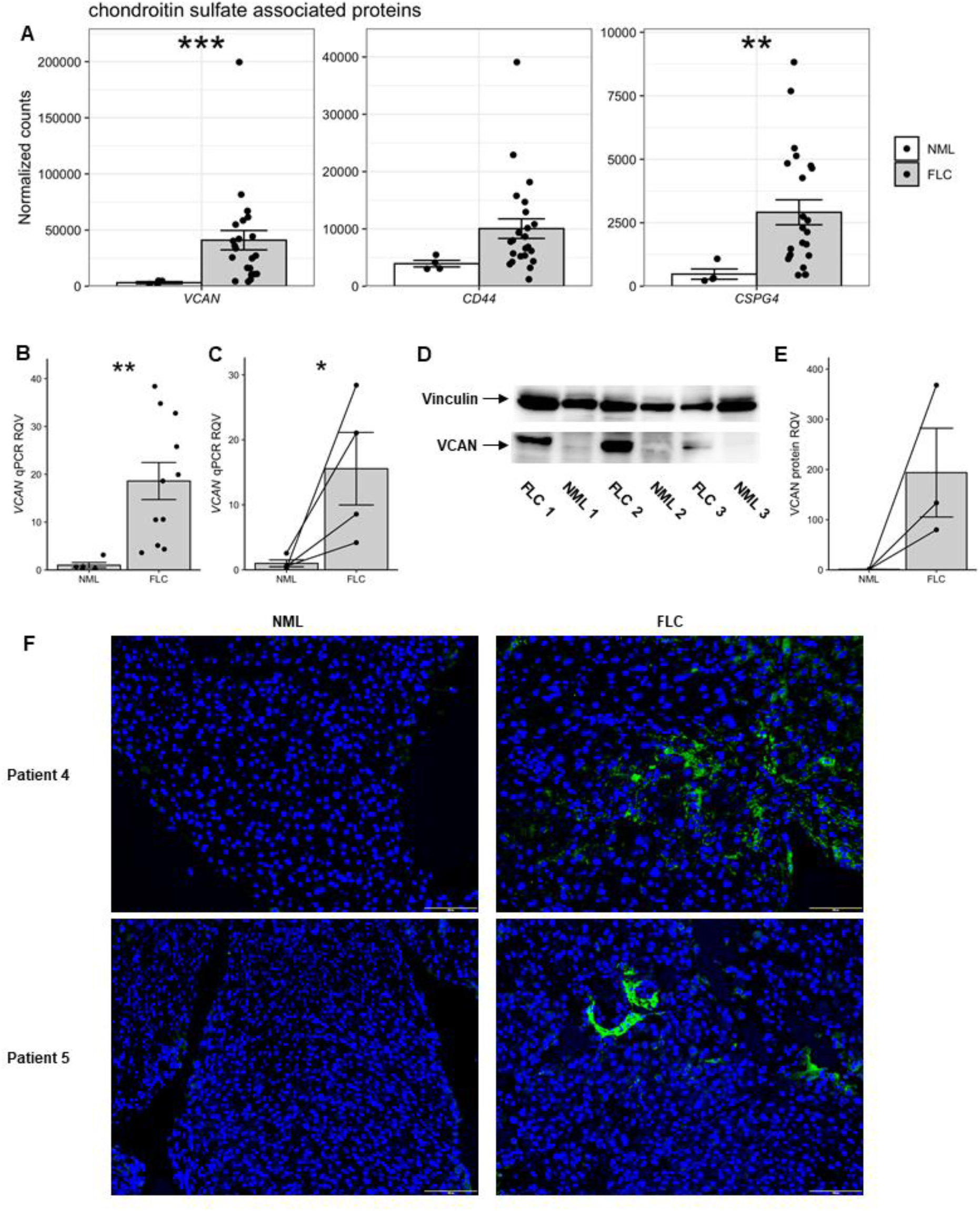
Chondroitin sulfate associated protein VCAN is aberrant in FLC. **(A)** Differential expression of *VCAN, CD44*, and *CSPG4* is shown as normalized counts in FLC (n=23) and NML (n=4). **(B)** Quantitative PCR showing the relative quantitative value (RQV) of VCAN in a separate cohort of FLC samples (n=11) compared to NML samples (n=5). **(C)** Quantitative PCR showing the relative quantitative value (RQV) of *VCAN* in a subset of FLC samples that have matched NML tissue (n=4). The matched NML/FLC samples are indicated with a line linking the two data points. **(D)** Immunoblot in matched FLC/NML samples (n=3) showing VCAN in the lower and vinculin in the upper panel. **(E)** VCAN protein levels normalized to vinculin levels are shown as relative quantitative value (RQV). The matched NML/FLC samples are indicated with a line linking the two data points. **(F)** IHF of VCAN protein in two independent patient-matched tissue samples is shown in green. Nuclei are blue. Scale bar equals 100 µm. \**P*<0.05, \*\**P*<0.01, \*\*\**P*<0.001, two-tailed Student’s t-test.

### Chondroitin sulfate GalNAc transferase 1 and versican are more altered in FLC than in most other cancer types and correlate with DNAJB1-PRKACA levels

Next, we sought to compare the expression of *CSGALNACT1* and *VCAN* in FLC to other cancer types. Specifically, we queried The Cancer Genome Atlas (TCGA) database, which houses RNA-seq data from 25 other cancer types. We found that the change in expression of *CSGALNACT1* in FLC (relative to corresponding non-malignant tissue) is second only to cholangiocarcinoma (**Figure 5A**). Strikingly, the change in expression of *VCAN* is greatest in FLC, followed by cholangiocarcinoma (**Figure 5B**). Further analysis revealed that *VCAN* and *CSGALNACT1* are correlated by RNA-seq (**Figure 5C**) as well as by RT-qPCR in an independent cohort (**Figure 5D**). Additionally, we found by RT-qPCR that the expression of DP correlates with both *VCAN* (**Figure 5E**) and *CSGALNACT1* (**Figure 5F**).

**Figure 5.**
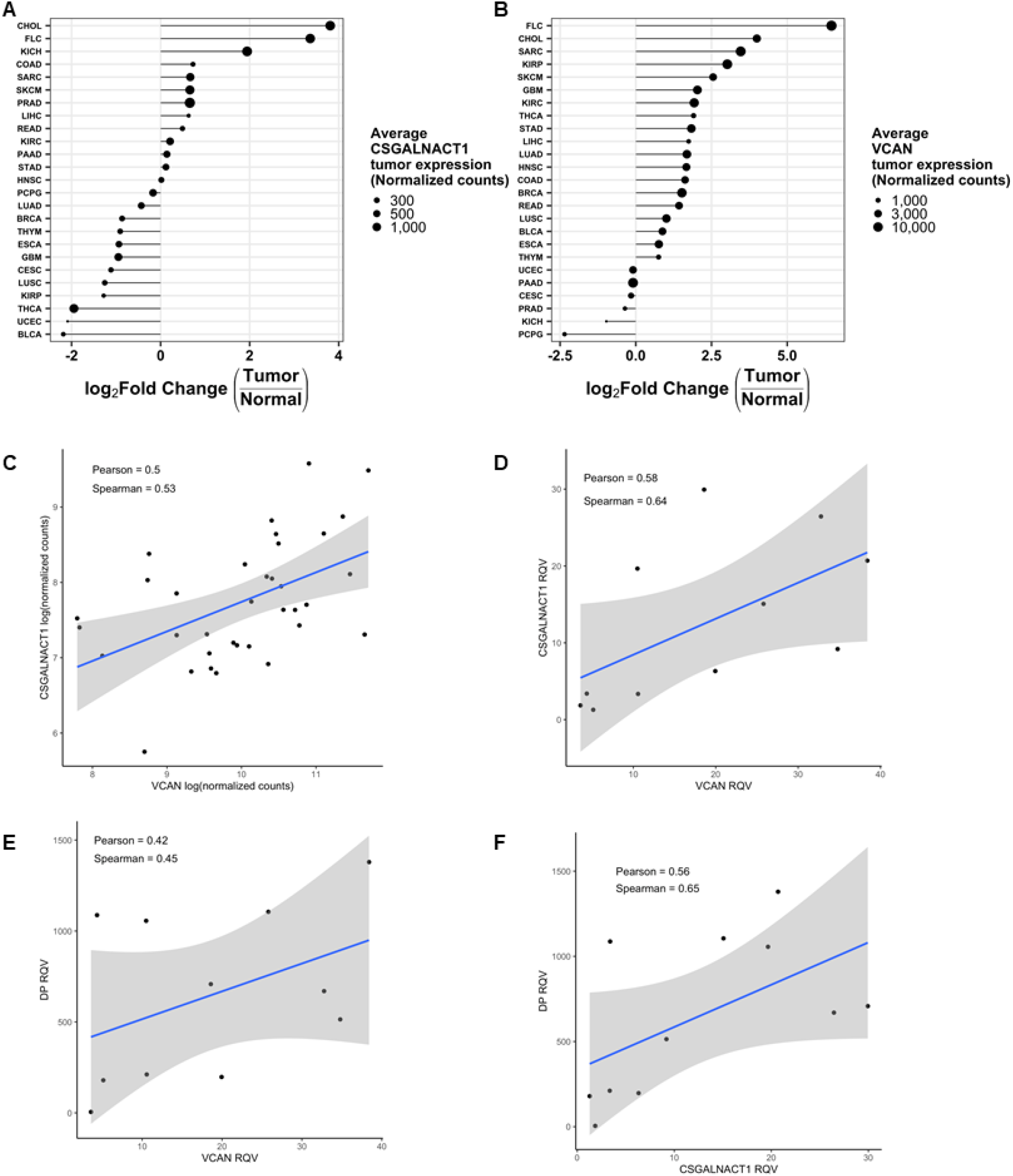
CSGALNACT1 and VCAN are more highly up-regulated in FLC than in almost all other cancers and correlate with DNAJB1-PRKACA levels. **(A)** Normalized counts of *CSGALNACT1* expression in RNA-seq data sets available for 25 tumor types in The Cancer Genome Atlas (TCGA). **(B)** Normalized counts of VCAN expression in RNA-seq data sets available for 25 tumor types in The Cancer Genome Atlas (TCGA). **(C)** Correlation between *CSGALNACT1* (y-axis) and *VCAN* (x-axis) shown as the log of normalized counts in FLC samples (n=27). **(D)** Correlation between *CSGALNACT1* (y-axis) and *VCAN* (x-axis) shown as the relative quantitative values from qPCR in a separate cohort of FLC samples (n=11). **(E)** Correlation between *DNAJB1-PRKACA* (y-axis) and *VCAN* (x-axis) shown as the relative quantitative values from RT-qPCR in FLC samples (n=11). **(F)** Correlation between *DNAJB1-PRKACA* (y-axis) to *CSGALNACT1* (x-axis) shown as the relative quantitative values from RT-qPCR in FLC samples (n=11). ACC, adenoid cystic carcinoma; BLCA, bladder urothelial carcinoma; BRCA, breast invasive carcinoma; CESC, cervical squamous cell and endocervical adenocarcinoma; CHOL, cholangiocarcinoma; COAD, colon adenocarcinoma; DLBC, diffuse large B-cell lymphoma; ESCA, esophageal carcinoma; FCL, fibrolamellar carcinoma samples analyzed in this study; GBM, glioblastoma; HNSC, head and neck squamous cell carcinoma; KICH, kidney chromophobe; KIRC, kidney renal papillary cell carcinoma; KIRP, kidney renal clear cell carcinoma; LAML, acute myeloid leukemia; LGG, lower grade glioma; LIHC, liver hepatocellular carcinoma; LUAD, lung adenocarcinoma; LUSC, lung squamous cell carcinoma; MESO, mesothelioma; PAAD, pancreatic adenocarcinoma; PCPG, pheochromocytoma and paraganglioma; PRAD, prostate adenocarcinoma; READ, rectum adenocarcinoma; SARC, sarcoma; SKCM, skin cutaneous melanoma; STAD, stomach adenocarcinoma; TGCT, testicular germ cell tumor; THCA, thyroid carcinoma; THYM, thymoma; UCEC, uterine corpus endometrial carcinoma; UCS, uterine carcinosarcoma; UVM, uveal melanoma.

### Versican is expressed in FLC transformed epithelial and tumor-associated, activated stellate cells in fibrolamellar carcinoma

To identify which cells are likely responsible for VCAN production and secretion we performed single-cell assay for transposase-accessible chromatin followed by sequencing (scATAC-seq) on NML, primary FLC tumor, and metastatic FLC tumor samples (n=3). We used a previously described nuclei isolation protocol ^(44)^ and obtained data on nearly 9,500 nuclei total. After data analysis with ArchR ^(45)^ (Methods), non-linear dimensionality reduction via UMAP revealed eight different clusters (**Figure 6A**). By analyzing open chromatin signal at established markers of human liver cell types ^(46)^, we assigned each cluster to a specific cell type (**Figure 6A, Supplemental Figure 4A-D**). We also analyzed open chromatin in the deleted region of chromosome 19 to identify the cells that likely harbor the deletion and therefore the DP fusion (**Supplemental Figure 5A, B**). We then queried for ATAC signal associated with *CSGALNACT1* and detected robust enrichment in the FLC primary and metastatic tumor transformed epithelial cell clusters (**Figure 6B**). As expected, there is little to no signal for open chromatin at *CSGALNACT1* in any non-malignant cell types (**Figure 6B, C**). A similar analysis for *VCAN* revealed the strongest signal in tumor-associated activated stellate cells and second-most in tumor transformed epithelial cells (**Figure 6D, E**). These findings suggest that while CS synthesis (via CSGALNACT1) is likely exclusively taking place in tumor transformed epithelial cells, the primary CS associated protein in FLC, VCAN, is produced and secreted from both activated stellate cells as well as tumor transformed epithelial cells (**Figure 7**).

**Figure 6.**
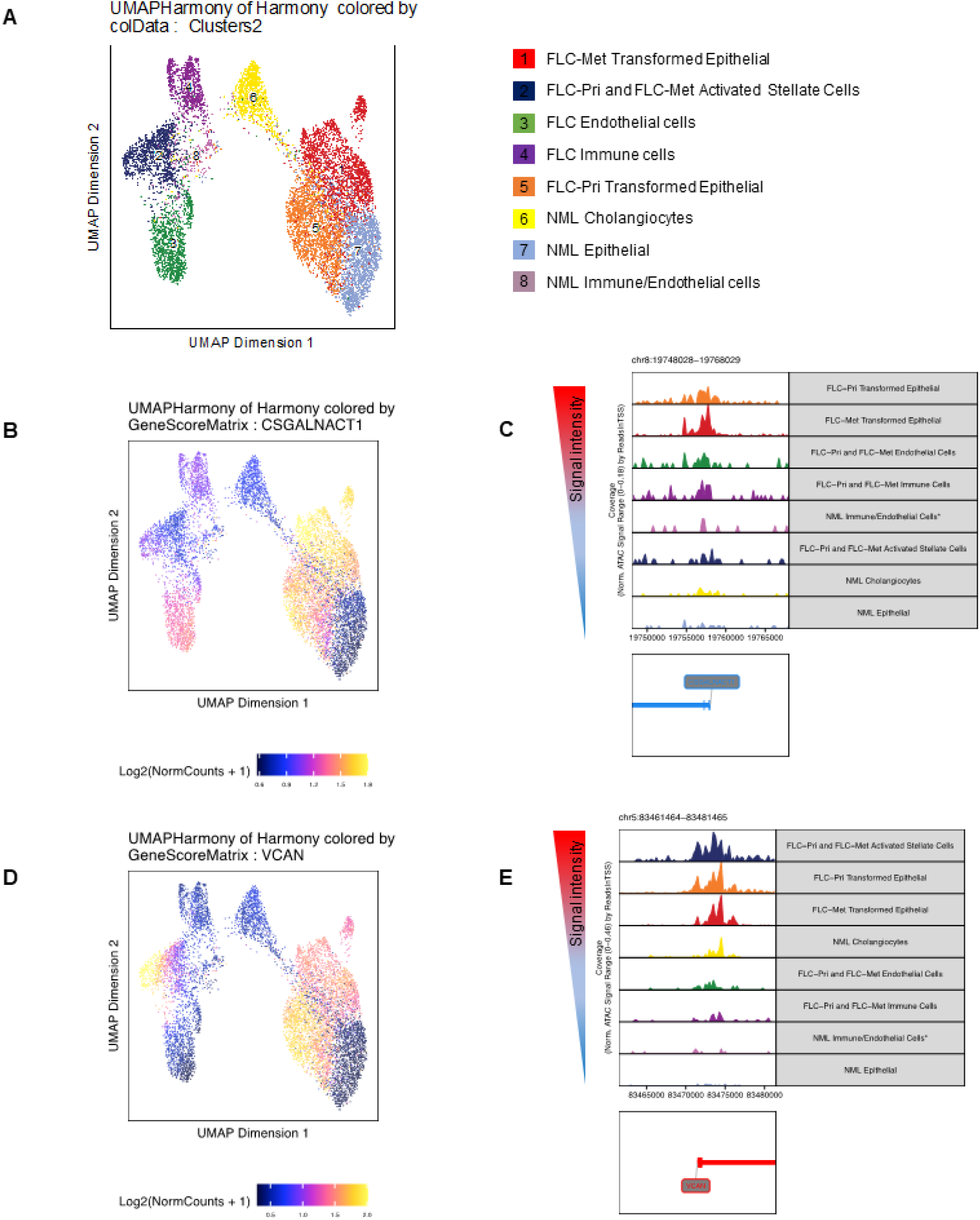
Activity at the *VCAN* locus is high in both tumor epithelial and activated stellate cells in FLC. **(A)** UMAP dimensional reduction showing eight cell clusters found in primary FLC, metastatic FLC, and NML tissue (∼9,500 nuclei). **(B)** Single-cell analysis of chromatin accessibility near the *CSGALNACT1* locus. Increasing signal is indicated by the color gradient (maximum signal is yellow and minimal signal is dark blue). **(C)** Genome tracks showing the location of open chromatin near the *CSGALNACT1* locus in each cell type. The annotated transcriptional start site for *CSGALNACT1* is shown at the bottom of the panel. **(D)** Single-cell analysis of chromatin accessibility near the *VCAN* locus. Increasing signal is indicated by the color gradient (maximum signal is yellow and the minimal signal is dark blue). **(E)** Genome tracks showing the location of open chromatin signal near the *VCAN* locus in each cell type. The annotated transcriptional start site for *VCAN* is shown at the bottom of the panel. Cell clusters are denoted through color-coding: NML hepatocytes in light blue, FLC primary tumor transformed epithelial cells in orange, FLC metastatic tumor transformed epithelial cells in red, NML cholangiocytes in yellow, FLC primary and metastatic activated stellate cells in dark blue, FLC primary and metastatic immune cells in dark purple, FLC primary and metastatic endothelial cells in green, and NML immune and endothelial cells in light purple.

**Figure 7.**
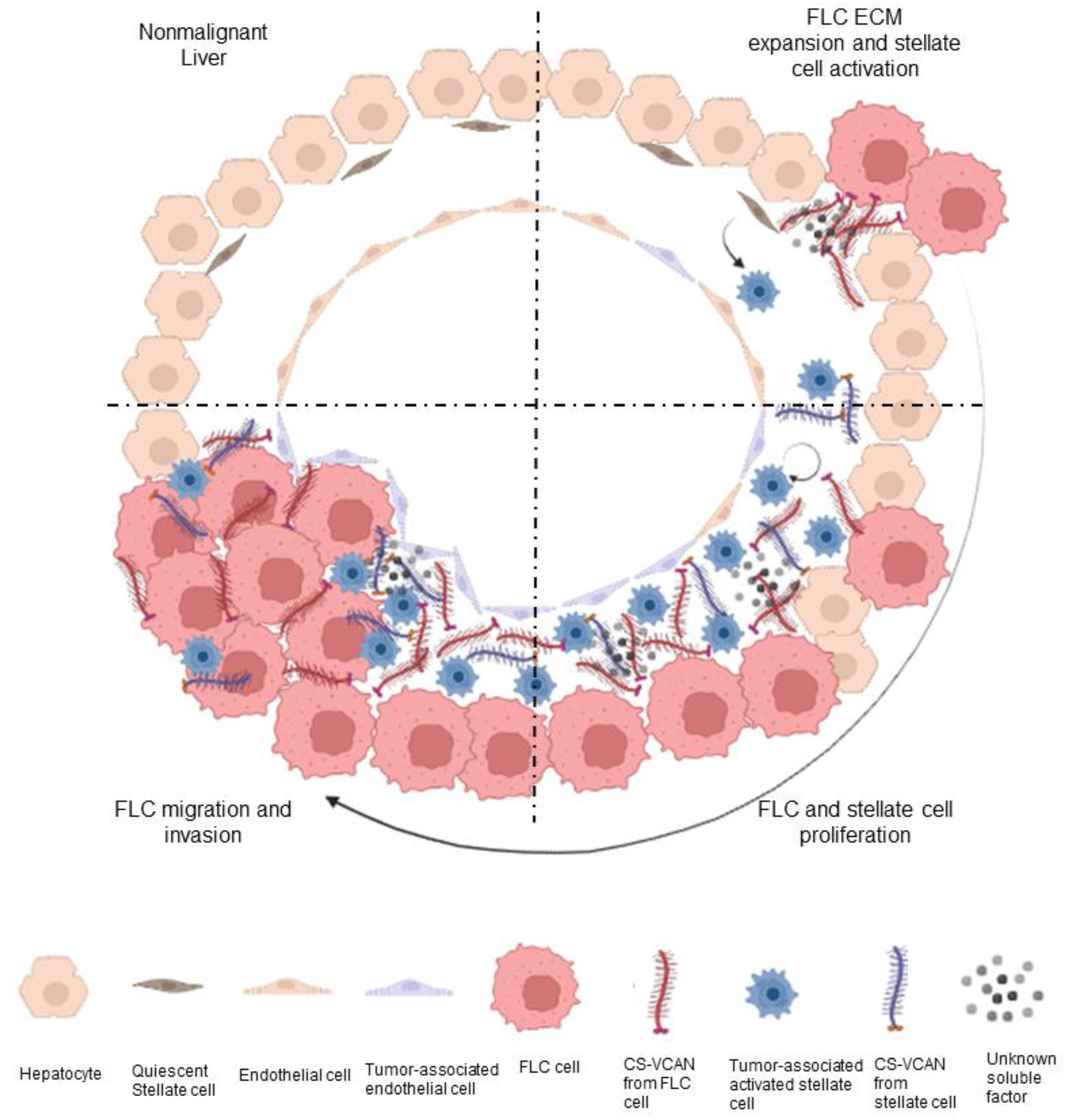
Model for the relevance of chondroitin sulfate and versican in FLC. A schematic showing the behaviors of non-malignant hepatocytes, quiescent and activated stellate cells, and endothelial cells with respect to chondroitin sulfate and versican during FLC progression.

## Discussion

FLC is an aggressive liver cancer that lacks an effective chemotherapeutic remedy. There are several factors contributing to low survival rates in FLC, including vague manifestations, lack of comorbidities, and resistance to general therapeutics. FLC is genetically characterized by the *DNAJB1-PRKACA* (DP) fusion, but efforts to identify specific inhibitors of DP, without targeting wild-type *PRKACA*, have been unsuccessful. Additionally, while it is known that DP is sufficient for tumor initiation, it is unclear whether DP expression is essential for tumor maintenance, progression, and metastasis. It has been established in the study of other cancer types that the pericellular environment, including proteoglycans, plays an important role in defining tumor behavior ^(12, 15)^. However, to date, no study has investigated GAGs and proteoglycans in FLC. In this study, we sought to bridge this knowledge gap.

The three major classes of GAGs are hyaluronic acid (HA), heparan sulfate (HS), and chondroitin sulfate (CS). Of these, only HA is found as a free polymer generated by hyaluronic acid synthetase 1-3 (HAS1-3). The expression levels of these enzymes were found to be extremely low in FLC and, therefore, they were not considered further. HS and CS chains are conjugated to proteins by a shared tetrasaccharide linker. The enzymes responsible for the synthesis of this linker exhibit robust expression in FLC but are unchanged relative to non-malignant liver (NML) tissue. The decision by cells to generate HS or CS sidechains is dependent on stoichiometric competition between the enzymes *N*-acetylglucosaminyltransferase (*EXTL2*), responsible for HS chains, and chondroitin GalNAc transferase (*CSGALNACT1*), responsible for CS chains. We found that the ratio of *CSGALNACT1* to *EXTL2* levels is dramatically elevated in FLC, pointing to CS chains as the key component of the extracellular matrix in FLC.

A major innovation and strength of this study is the development and implementation of a highly sensitive method for quantifying HS and CS disaccharides in patient tissues. This novel assay confirmed that the alterations in expression of CS biosynthetic genes observed in FLC lead to dramatic changes in CS abundance. The quantity of CS in FLC tissue relative to NML was greater than 5.9-fold and the relative difference between CS and HS in tumor tissue was 4.3-fold. Additionally, this assay independently quantifies sulfated forms of CS and HS disaccharides. As HS abundance is unchanged in FLC tissue, it is not surprising that there are no significant differences in the seven sulfated forms of HS that we measured. The analysis of sulfated forms of CS showed that nonsulfated CS and two forms of monosulfated CS (CS di-4S and CS di-6S) are significantly increased in FLC tissue. The increased abundance of CS di-4S and CS di-6S is concordant with gene expression increases in associated chondroitin sulfotransferases (*CHST11/3*), again showing that changes in gene expression accurately correspond with changes in chemical abundance. It has been observed in hepatocellular carcinoma (HCC) that increased expression of *CHST11/13*, which are functionally equivalent, are upregulated in metastatic samples ^(47)^, which may promote sustained Wnt signaling ^(48)^. One limitation of the quantitative analysis for the CS chains with di-2S4S should be noted. We were unable to synthesize ^13^C-labeled di2S4S; therefore, the quantitation was completed using the relative quantitation method. However, this should not affect our conclusions, as the levels of di-2S4S are the same in FLC compared to NML tissues.

Given the abundant increase of CS in FLC tissue, and the high expression of the CS-associated protein VCAN, we conjectured that the levels of *CSGALNACT1* and *VCAN* would be correlated. Indeed, we found that there is a positive correlation between the two genes, and between either gene and DP. Consistent with this finding, in a previous report we had demonstrated that the expression of DP in an FLC cell model increases VCAN expression ^(5)^. We found in this study that the levels of VCAN are upregulated in FLC more than in any other cancer type for which expression data is publicly available through TCGA. Cholangiocarcinoma (CCA) is the closest to FLC in terms of *CSGALNACT1* and *VCAN* upregulation. Intriguingly, chondroitin polymerizing factor (*CHPF*) has been reported recently to promote CCA cell growth and invasive potential ^(49)^. Additionally, chondroitin synthase 1 (*CHSY1*) has been reported to suppress apoptosis in colorectal cancer ^(50)^ and promote migration in hepatocellular carcinoma (HCC) ^(51)^.

Another major component of this study is the first-ever single-cell ATAC-seq analysis of FLC. All prior genome-scale analyses of FLC have been performed on bulk tumors and the only published single-cell analysis related to this cancer involves a patient-derived xenograft (PDX), not a primary tumor ^(52)^. Our study overcomes these limitations and provides the first glimpse into FLC tumor tissue complexity. We focused on resolving gene locus activity at single-cell resolution and identified chromatin accessibility at the *CSGALNACT1* and *VCAN* loci in the primary population of FLC cells. Transformed epithelial cells were identified as the primary source of *CSGLANACT1* locus activity, whereas proliferating stellate cells were found to be the cell type with the strongest *VCAN* signal. This finding suggests that communication between transformed DP+ liver epithelial cells and stellate cells may be critical to FLC disease progression.

The expression and secretion of VCAN from activated stellate cells is a normal response to liver injury ^(53-56)^. The data generated in this study suggest that FLC cells upregulate *CSGALNACT1* and *VCAN* in a DP-dependent manner and begin secreting CS-VCAN proteoglycan into the extracellular matrix. The increased accumulation of VCAN may sequester higher concentrations of growth factors, or increase mechanical tension, and induce the activation of quiescent stellate cells. Upon activation, these stellate cells proliferate and secrete VCAN as a normal response to a perceived injury. However, the rapid accumulation of VCAN may promote proliferation and migration of FLC cells, creating a feed-forward loop, directly though CS signaling, mechano-transduction, and/or growth factor presentation.

The role for stellate cells to promote fibrosis and predispose the liver to cancer formation is well established ^(57)^ and multiple HS proteoglycans have been implicated in this role, including syndecans ^(58-61)^, glypicans ^(62-64)^, and even free HS disaccharides ^(65)^. Additionally, CS proteoglycans, including VCAN and CD44, have been implicated in both hepatic fibrosis and HCC formation ^(31, 66, 67)^. However, these findings suggest that stellate-mediated fibrosis precedes and contributes to cancer formation. Given that FLC patients lack underlying fibrotic conditions, such as cirrhosis, the relationship in FLC may be fundamentally different. Further investigation in the transcriptional activation of *VCAN* in FLC and the identification of VCAN associated growth factors is warranted.

We have implemented several novel methods to provide the first high-resolution analysis of proteoglycan biology in FLC tumors. Our findings motivate further investigation of VCAN in FLC progression. Future research may also study the effects of VCAN inhibitors on FLC cell drug resistance, growth and/or metastasis.

## Acknowledgments

We are very grateful for funding from the Fibrolamellar Cancer Foundation (awarded to P.S.), which permitted the execution of this project. We acknowledge the service provided by the Cornell Genomics Facility for sequencing. We are especially indebted to the Cornell Genomics Innovation hub, including Jen Grenier, Adrian McNairn, and Paul Munn, for their invaluable insight and input for single-cell ATAC library preparation. Finally, we also thank Larry Bonassar, Jongkil Kim, and Cora Demler for valuable discussions during the study. Chemical structures were created with Chemdraw (by PerkinElmer). Schematics were created with BioRender.com.

## Author contributions

Conceptualization, A.B.F., Jian.L., P.S.; Methodology, Jine.L., Jian.L., B.S.; Software, A.R.F., M.K.; Formal Analysis, A.B.F., Jine.L., A.R.F., M.K.; Investigation, A.B.F., Jine.L., A.R.F.; Data Curation, A.R.F., M.K.; Writing – Original Draft, A.B.F., Jian.L., P.S.; Writing – Reviewing & Editing, Jine.L., A.R.F., M.K., P.D.S, Z.W., L.M.R.; Visualization, A.B.F., Jine.L., A.R.F., M.K.; Funding Acquisition, P.S.; Resources, P.D.S, Z.W.; Supervision, Jian.L., P.S.

## Inclusion and Diversity

We worked to ensure gender balance in the recruitment of human subjects. We worked to ensure ethnic or other types of diversity in the recruitment of human subjects. One or more of the authors of this paper self-identifies as an underrepresented ethnic minority in science. While citing references scientifically relevant for this work, we also actively worked to promote gender balance in our reference list. The author list of this paper includes contributors from the location where the research was conducted who participated in the data collection, design, analysis, and/or interpretation of the work.

## Figure Legends

**Supplemental Figure 1.**
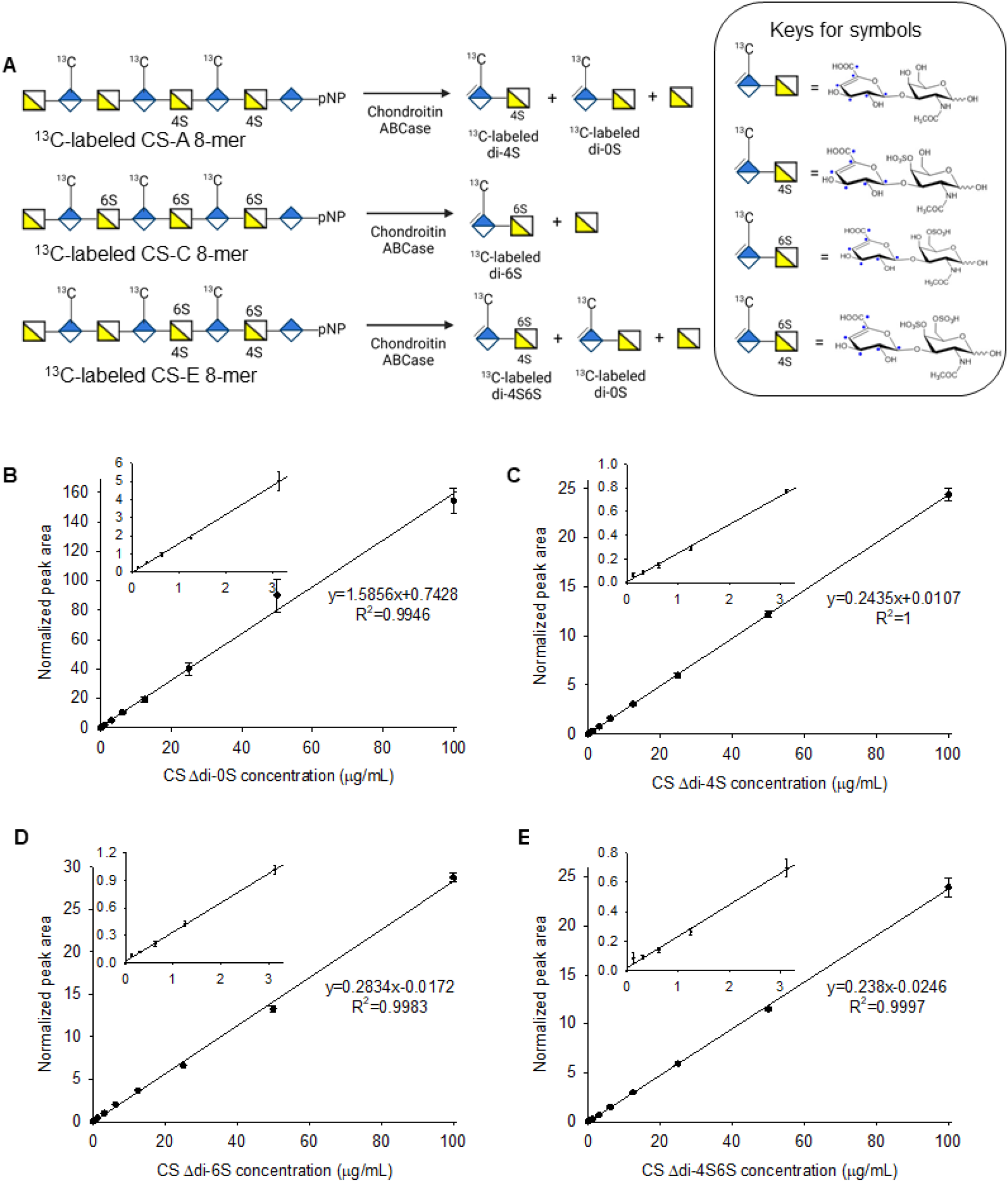
Quantification of chondroitin sulfate disaccharides. **(A)** Schematic of preparation of ^13^C-labeled CS disaccharides. Four ^13^C-labeled CS disaccharide calibrants were prepared from chondroitin ABCase digestion of three ^13^C-labeled 8-mers. The disaccharides were purified to homogeneity after a Q-Sepharose column purification. The ^13^C-labeled carbon atoms in the DUA residue are indicated with blue dots. pNP represents p-nitrophenyl. **(B)** The curve and linear equation of normalized peak area as a function of concentration for Δdi-0S are shown. The concentration of Δdi-0S used for LC-MS/MS analysis were 0.125, 0.313, 0.625, 1.25, 3.13, 6.25, 12.5, 25, 50 and 100 μg/mL, mixing with 0.8 μg/mL disaccharide calibrant Δdi-4S. Data represent means ± S.D. (n=3) **(C)** The curve and linear equation of normalized peak area as a function of concentration for Δdi-4S are shown. The concentration of Δdi-4S used for LC-MS/MS analysis were 0.125, 0.313, 0.625, 1.25, 3.13, 6.25, 12.5, 25, 50 and 100 μg/mL, mixing with 4 μg/mL disaccharide calibrant Δdi-4S. Data represent means ± S.D. (n=3) **(D)** The curve and linear equation of normalized peak area as a function of concentration for Δdi-6S are shown. The concentration of Δdi-6S used for LC-MS/MS analysis were 0.125, 0.313, 0.625, 1.25, 3.13, 6.25, 12.5, 25, 50 and 100 μg/mL, mixing with 4 μg/mL disaccharide calibrant Δdi-6S. Data represent means ± S.D. (n=3) **(E)** The curve and linear equation of normalized peak area as a function of concentration for Δdi-4S6S are shown. The concentration of Δdi-4S6S used for LC-MS/MS analysis were 0.125, 0.313, 0.625, 1.25, 3.13, 6.25, 12.5, 25, 50 and 100 μg/mL, mixing with 4 μg/mL disaccharide calibrant Δdi-4S6S. Data represent means ± S.D. (n=3)

**Supplemental Figure 2.**
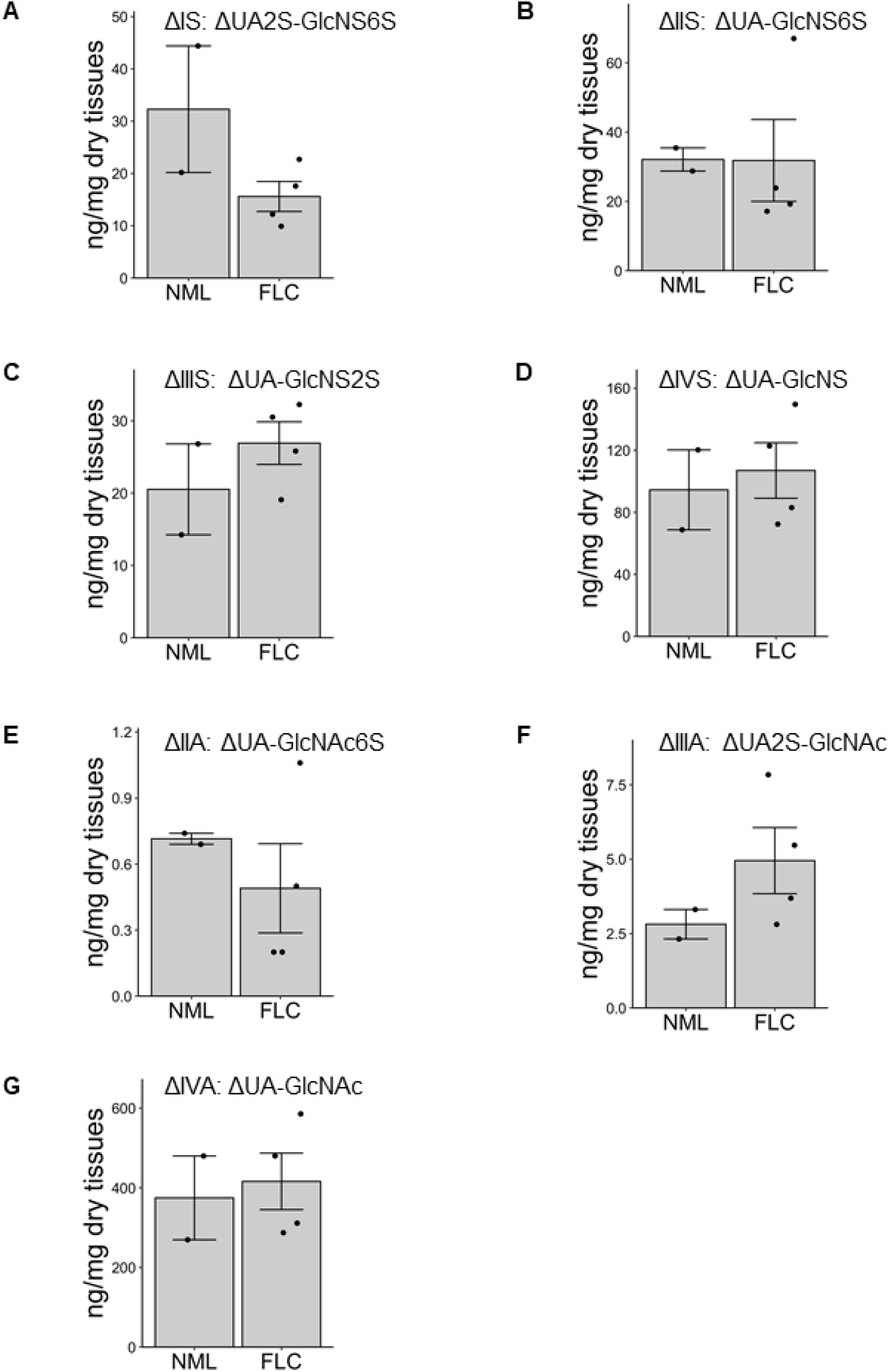
Quantification of heparan sulfate disaccharides. **(A)** Nanograms of HS ΔUA2S-GlcNS6S (ΔIS) per milligram of dry tissue in FLC (n=4) and NML (n=2) tissue. **(B)** Nanograms of HS ΔUA-GlcNS6S (ΔIIS) per milligram of dry tissue in FLC (n=4) and NML (n=2) tissue. **(C)** Nanograms of HS ΔUA-GlcNS (ΔIIIS) per milligram of dry tissue in FLC (n=4) and NML (n=2) tissue. **(D)** Nanograms of HS ΔUA-GlcNS (ΔIVS) per milligram of dry tissue in FLC (n=4) and NML (n=2) tissue. **(E)** Nanograms of HS ΔUA-GlcNAc6S (ΔIIA) per milligram of dry tissue in FLC (n=4) and NML (n=2) tissue. **(F)** Nanograms of HS ΔUA2S-GlcNAc (ΔIIIA) per milligram of dry tissue in FLC (n=4) and NML (n=2) tissue. **(G)** Nanograms of HS ΔUA-GlcNAc (ΔIVA) per milligram of dry tissue in FLC (n=4) and NML (n=2) tissue.

**Supplemental Figure 3.**
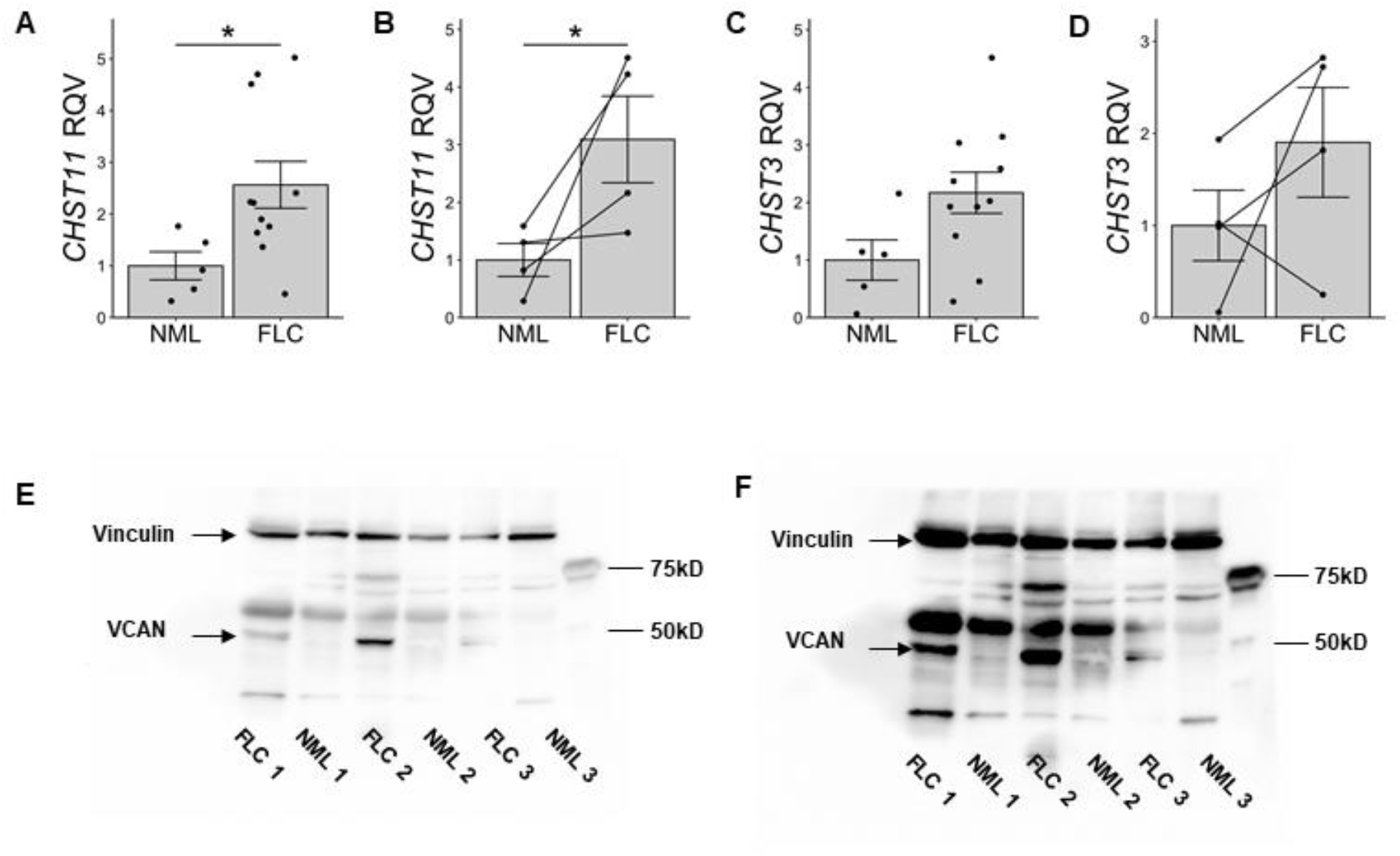
Additional gene expression analysis in FLC samples and additional VCAN immunoblots. **(A)** Quantitative PCR showing the relative quantitative value (RQV) of *CHST11* in a subset of FLC samples that have matched NML tissue (n=4). **(B)** The matched NML/FLC samples are indicated with a line linking the two data points. **(C)** Quantitative PCR showing the relative quantitative value (RQV) of *CHST3* in a subset of FLC samples included in this study (n=11) compared to NML samples (n=5). **(D)** Quantitative PCR showing the relative quantitative value (RQV) of *CHST3* in a subset of FLC samples that have matched NML tissue (n=4). The matched NML/FLC samples are indicated with a line linking the two data points. **(E)** Short and **(F)** long exposure VCAN immunoblot from Figure 4D shown in entirety. Vinculin and versican protein are denoted with arrows and molecular weight standards are denoted by dashes. Samples are labeled at the bottom of the panel. The short exposure blot was used for quantification in Figure 4E. \**P*<0.05, two-tailed Student’s t-test.

**Supplemental Figure 4.**
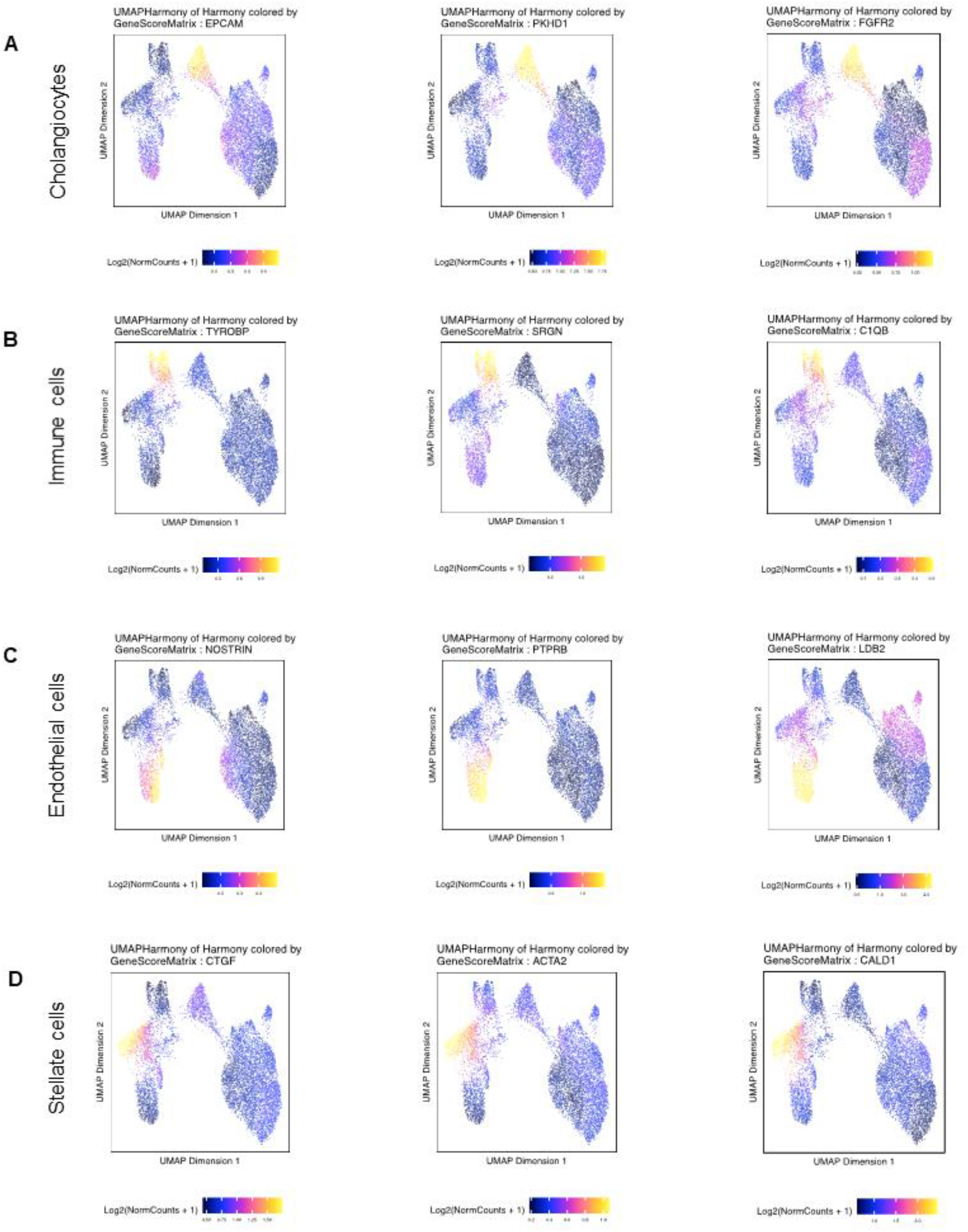
Marker genes of cell clusters from the single-cell ATAC data. **(A)** Signal intensity of chromatin accessibility near the *EPCAM, PKHD1*, and FGFR2 gene loci denoting cholangiocytes. **(B)** Signal intensity of chromatin accessibility near the *TYROBP, SRGN*, and *C1QB* gene loci denoting immune cells. **(C)** Signal intensity of chromatin accessibility near the *NOSTRIN, PTPRB*, and *LDB2* gene loci denoting endothelial cells. **(D)** Signal intensity of chromatin accessibility near the *CTGF, ACTA2*, and *CALD1* gene loci denoting stellate cells. Increasing signal is indicated by the color gradient (maximum signal is yellow and minimal signal is dark blue).

**Supplemental Figure 5.**
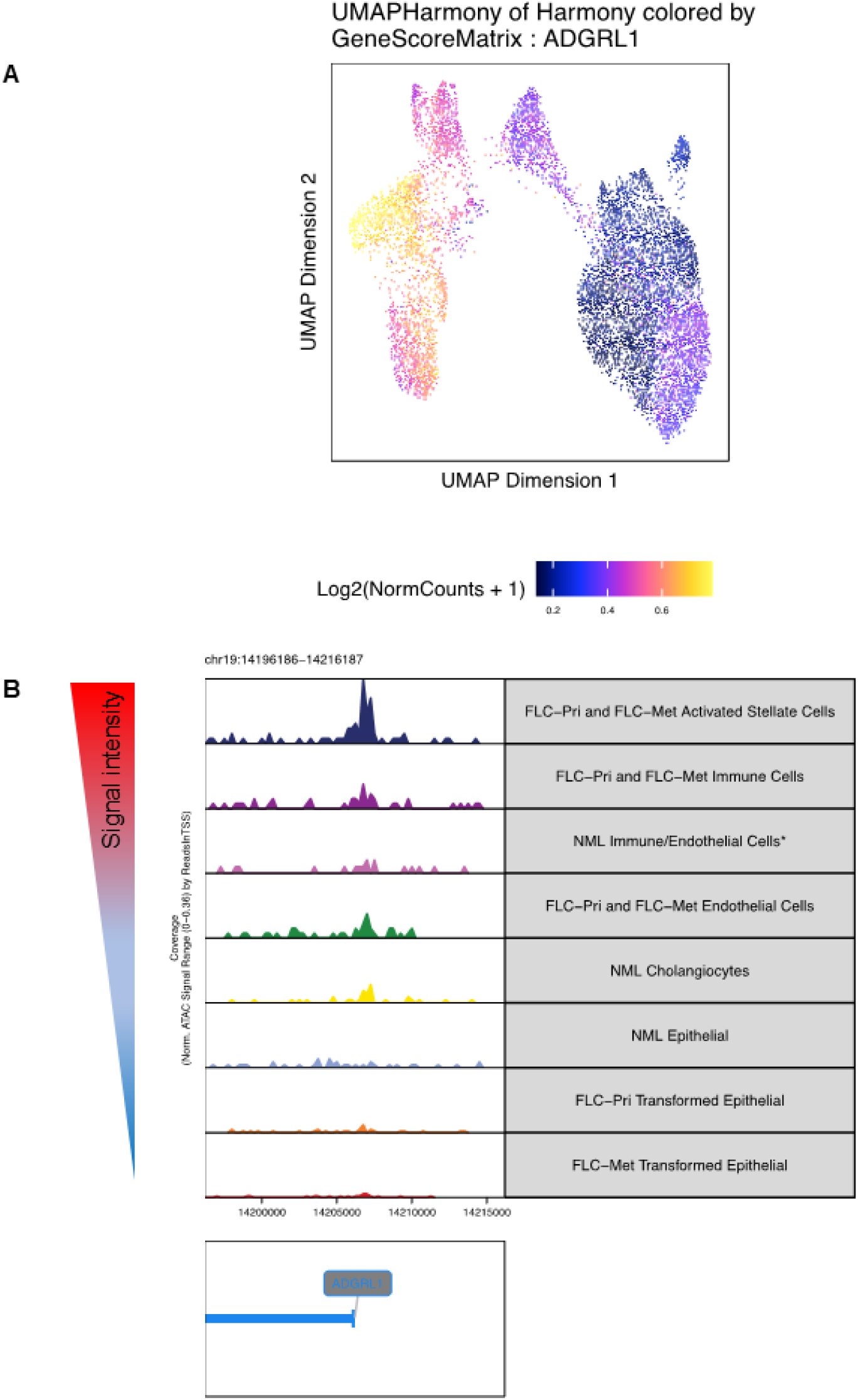
Identification of cells that are likely positive for the signature deletion event in FLC. **(A)** Signal intensity of chromatin accessibility near the *ADGRL1* gene locus. **(B)** Genome tracks showing the location of open chromatin signal near the *ADGRL1* locus in each cell type. The annotated transcriptional start site for *ADGRL1* is shown at the bottom of the panel.

